# Functional analysis of conserved *C. elegans* bHLH family members uncovers lifespan control by a peptidergic hub neuron

**DOI:** 10.1101/2024.07.12.603289

**Authors:** G. Robert Aguilar, Berta Vidal, Hongzhu Ji, Joke Evenblij, Hongfei Ji, Giulio Valperga, Chien-Po Liao, Christopher Fang-Yen, Oliver Hobert

## Abstract

Throughout the animal kingdom, several members of the basic helix-loop-helix (bHLH) family act as proneural genes during early steps of nervous system development. Roles of bHLH genes in specifying terminal differentiation of postmitotic neurons have been less extensively studied. We analyze here the function of five *C. elegans* bHLH genes, falling into three phylogenetically conserved subfamilies, which are continuously expressed in a very small number of postmitotic neurons in the central nervous system. We show (a) that two orthologs of the vertebrate bHLHb4/b5 genes, called *hlh-17* and *hlh-32,* function redundantly to specify the identity of a single head interneuron (AUA), as well as an individual motor neuron (VB2), (b) that the *PTF1a* ortholog *hlh-13* acts as a terminal selector to control terminal differentiation and function of the sole octopaminergic neuron class in *C. elegans*, RIC, and (c) that the NHLH1/2 ortholog *hlh-15* controls terminal differentiation and function of the peptidergic AVK head interneuron class, a known neuropeptidergic signaling hub in the animal. Strikingly, through null mutant analysis and cell-specific rescue experiments, we find that loss of *hlh-15/NHLH* in the peptidergic AVK neurons and the resulting abrogation of neuropeptide secretion causes a substantially expanded lifespan of the animal, revealing an unanticipated impact of a central, peptidergic hub neuron in regulating lifespan, which we propose to be akin to hypothalamic control of lifespan in vertebrates. Taken together, our functional analysis reveals themes of bHLH gene function during terminal differentiation that are complementary to the earlier lineage specification roles of other bHLH family members. However, such late functions are much more sparsely employed by members of the bHLH transcription factor family, compared to the function of the much more broadly employed homeodomain transcription factor family.

## INTRODUCTION

Basic helix-loop-helix (bHLH) transcription factors constitute a large, deeply conserved family of transcription factors with wide-ranging roles in animal development and physiology (reviewed in (Jan and Jan 1994; Hassan and Bellen 2000; Massari and Murre 2000; Bertrand *et al*. 2002; Wang and Baker 2015; Guillemot and Hassan 2017; Baker and Brown 2018)). bHLH proteins have been categorized into six higher order groups (Group A to F) based on intrinsic sequence features of the bHLH domain and/or the presence of additional domains (Ledent *et al*. 2002; Simionato *et al*. 2008; Gyoja 2014). Group A is the largest of these groups and contains proteins with deeply conserved developmental patterning functions in several tissue types, most prominently in the nervous system (e.g. Atonal, ASC families) and muscle (MyoD, Twist). Many members of this group heterodimerize with a common subunit, the E-proteins in vertebrates, Daughterless in *Drosophila* and HLH-2 in *C. elegans*, which are also all members of group A. This common subunit is often referred to as a “class I” bHLH protein, while other group A members that heterodimerize with the class I protein are referred to as “class II” bHLH proteins (Massari and Murre 2000). While members of all major bHLH groups (i.e. group A to F) appear to have been present in single cell organisms (Sebe-PEDROS *et al*. 2011), group A bHLH genes underwent major expansions during metazoan evolution (Gyoja 2014) and hence are possible contributors to animal cell type diversification.

The nematode *C. elegans* contains 20 group A family members, including orthologs of prominent developmental patterning genes, such as mesodermal patterning genes MyoD (*hlh-1* in *C. elegans*) and Twist (*hlh-8*) as well as orthologs of neuronal patterning genes Atonal (*lin-32*), Neurogenin (*ngn-1*), NeuroD (*cnd-1*) and others (Ledent *et al*. 2002; Simionato *et al*. 2008)(**Fig.1A**). As exemplified by our comprehensive analysis of homeobox gene expression and function (Reilly *et al*. 2020; Hobert 2021; Reilly *et al*. 2022), we reason that broad, family-wide, and animal-wide analyses of transcription factor families may reveal the presence (or absence) of common themes in their expression and function. Hence, over the past few years, our laboratory has engaged in a systematic analysis of several of the group A bHLH proteins throughout the entire *C. elegans* nervous system, revealing common and divergent themes in the function of class II proteins, including the ASC homologs *hlh-14* and *hlh-4,* the Atonal homolog *lin-32*, and the class I Daughterless homolog *hlh-2* (Poole *et al*. 2011; Masoudi *et al*. 2018; Masoudi *et al*. 2021; Masoudi *et al*. 2022).

**Fig. 1:**
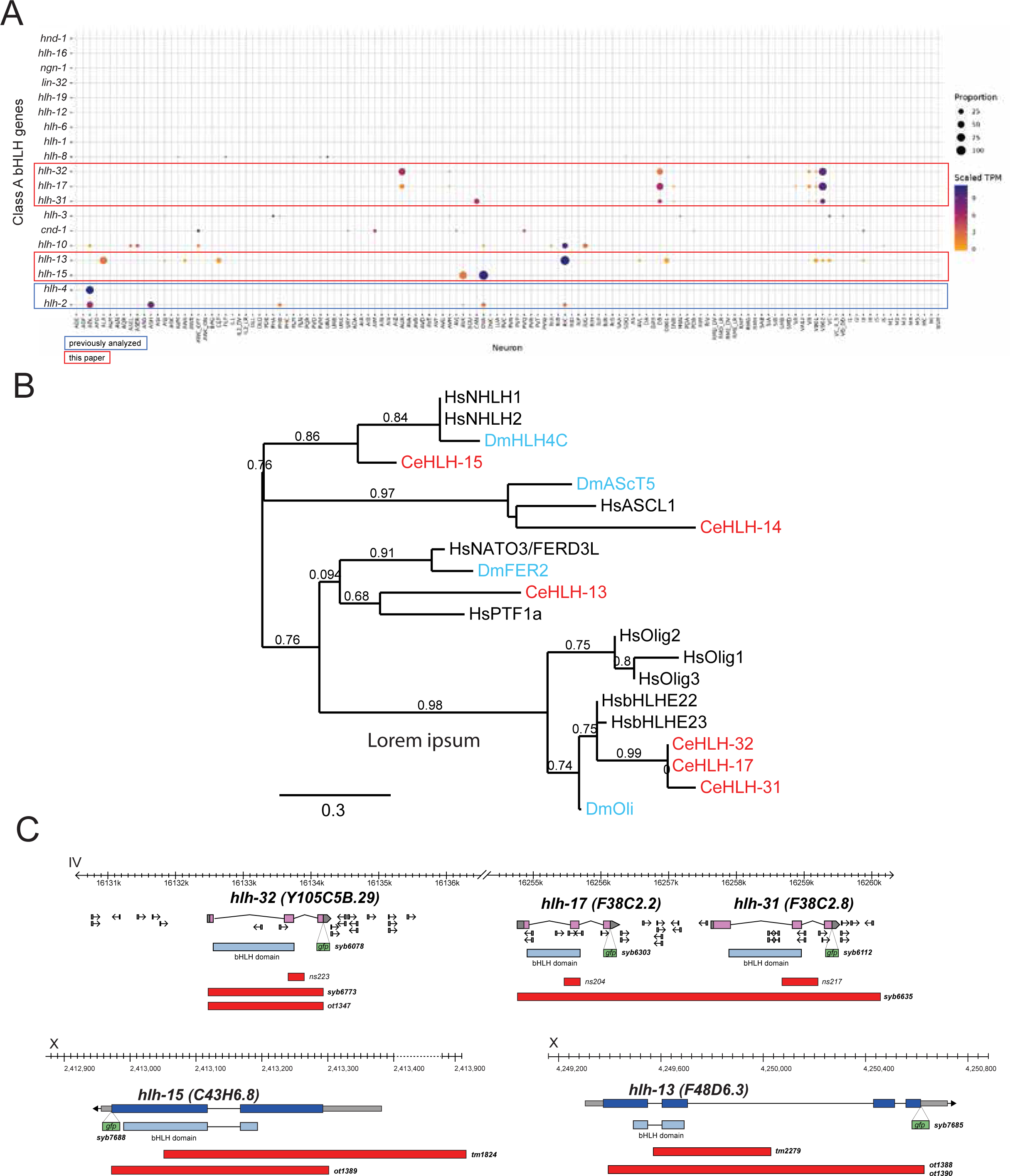
Group A class members in *C. elegans*. **A:** Expression of group A class members in *C. elegans* in the CeNGEN scRNAseq dataset (Taylor *et al*. 2021). Orthologs are as follows (Ledent *et al*. 2002; Simionato *et al*. 2008): *hlh-2 =* E12/47, D.m. Daughterless; *hlh-1=* MyoD; *hlh-3, 4, 6, 12, 19:* ASC-related; *lin-32 =* Atonal homolog; *cnd-1 =* NeuroD; *ngn-1 =* Neurogenin; *hlh-17, hlh-31, hlh-32 =* bHLHe4/5 orthologs (see **Fig.S1**); *hlh-16 =* Olig related (see **Fig.S1**); *hlh-10, hlh-12 =* TCF21 homolog ; *hlh-13 = PTF1a* homolog *; hlh-15 =* NHLH1 & 2 homolog; *hlh-8 =* Twist; *hnd-1 =* Hand. **B:** Phylogenetic relationship of the bHLH domain sequences of select group A bHLH family members. A representative ASc family member (HLH-14) and its orthologs are included. There are two additional FER paralogs in *Drosophila* not included in this tree. Phylogenetic trees were generated at phylogeny.fr (Dereeper *et al*. 2008) with default parameters. **C:** Locus with deletion and reporter alleles for all examined genes (*hlh-13, 15, 17, 31, 32*).

In this paper, we have used the recently released single cell RNA sequencing (scRNAseq) atlas of the entire, mature nervous system of *C. elegans* (Taylor et al. 2021) to choose for functional analysis a set of five postmitotically expressed, deeply conserved group A family members. These five bHLH proteins were chosen based on their very restricted, strong expression in individual, mature cell types of the nervous system, where their expression is maintained throughout the life of the animal, as indicated by recent scRNAseq analysis (**Fig.1A**)(Taylor *et al*. 2021). These proteins include the sole *C. elegans* ortholog of the vertebrate PTF1a and NATO3 genes, *hlh-13,* the sole *C. elegans* ortholog of the vertebrate NHLH1 and NHLH2 genes, *hlh-15* and the three *C. elegans* orthologs of the vertebrate bHLHb4/b5 genes, called *hlh-17, hlh-31* and *hlh-32* (**Fig.1B**). Each of these vertebrate proteins had been implicated in neuronal differentiation in select parts of the mammalian nervous system (Bramblett *et al*. 2004; Feng *et al*. 2006; Joshi *et al*. 2008; Ono *et al*. 2010; Ross *et al*. 2010; Skaggs *et al*. 2011; Ross *et al*. 2012; Good and Braun 2013; Huang *et al*. 2014; Jin and Xiang 2019).

We make use of the more compact nature of the *C. elegans* nervous system to garner a systems-wide view of the functions of their *C. elegans* orthologs and to more precisely assess their function using the many tools that *C. elegans* offers to dissect neuronal differentiation programs. Our analysis is a further stepping-stone in our attempt to generate an encyclopedic map of neuronal identity regulators in *C. elegans.* One emerging view of such encyclopedic analyses is that, compared to another prominent transcription factor family, the homeodomain family, bHLH protein function appears more sparsely employed during late differentiation steps in the nervous system.

## MATERIAL AND METHODS

### C. elegans strains

All strains used in this study are listed in **Table S1**. Strains were maintained as previously described (Brenner 1974).

### *C. elegans* genome engineered strains

Deletion alleles for *hlh-32(ot1347), hlh-13(ot1388), hlh-13(ot1390)* and *hlh-15(ot1389)* were generated using CRISPR/Cas9 genome engineering (Dokshin *et al*. 2018). Multiple alleles encompassing the same genomic regions, as shown in **Fig.1**, represent the same deletion generated in different backgrounds. Deletions were made using two crRNAs repaired with an ssODN, as described below:

*hlh-32(ot1347):* crRNAs (tttcagggtgtcgttagatt and catccgaaggagattagcgc), ssODN (cgctggtgatagattcctggatgttcaacaaattgcgctggtgatagattcctggatgttcaacaaattg)

*hlh-13(ot1388)* and *hlh-13(ot1390):* crRNAs (gaagctgtcatttacataag and acagagtttgtttaggcaat), ssODN (accaattacaattgtgaattcgagcagaaccacttGCCTAAacaaactctgtgtatgcgtaaacggcaga)

*hlh-15(ot1389):* crRNAs(ttgaatagtggaacacttgc and tgaattcacaaccttcacag), ssODN(atctgttactcgttttcctatcctctattccagcacagtggccaaccgatatatatatatatatagttcc)

A number of CRISPR/Cas9-engineered reporter and deletion strains were generated by SunyBiotech (indicated by the *syb* allele designation; see **Table S1**). These include C-terminal *gfp* reporter insertions for all the 5 bHLH genes examined here and SL2::gfp::H2B reporter cassette insertions at the C-terminus of genes that serve as cell fate markers and/or potential targets of bHLH genes (*nlp-49, kcc-3, col-105, pdf-1, flp-7, bcat-1*). All other previously published cell fate markers used here are also listed in **Table S1**.

### *C. elegans* transgenic strains

To establish the *sshk-1p::gfp* reporter transgene, a marker for AUA neurons, the 370bp intergenic region upstream of *sshk-1* was fused to *gfp* followed by the 3’ untranslated region of *unc-54,* as previously described (Hobert 2002). Primers were:

*sshk-1* promoter (370 bp): **fwd** *CTGTTGACTAATCTCACAGC* **rev** *AGGAAAACTTTCAAATGAGAGG.* The amplicon was injected together with a *pha-1* wildtype copy into in a *pha-1(e2123)* mutant strain to generate *otEx8199*.

To generate *flp-1p::genomic hlh-15::SL2::TagRFP*, a 0.4 kb *flp-1* promoter sequence from pCS169 (*flp-1p(trc)::ICE*) (Oranth *et al*. 2018) attached to the full *hlh-15* genomic sequence (*flp-1p::hlh-15 genomic fragment* containing exons and introns) was fully synthesized by Azenta Life Sciences and amplified using the following primers: **fwd** *GGAAATGAAATCAGGAAACAGCTATGACCATGAGCTTAATTCCTAAAAACCC* **rev** *GGTGAAAGTAGGATGAGACAGCCGGCCGTTATTGAAGCAAGTTGTCTAGAAAAC*.

This amplicon was then ligated to the SL2::TagRFP backbone amplified from pCPL11 (*srab-20p::lin-29a::SL2::TagRFP*) using Gibson cloning. *hlh-15(ot1389)* mutant animals were then injected with the following concentrations of constructs: *flp-1p::ghlh-15::SL2::tagRFP 25ng/ul, Pttx-3::GFP 25ng/ul, pBS 150ng/ul* to generate *otEx8247* and *otEx8248*.

All other previously described transgenic strains used in this study are listed in **Table S1.**

### Automated worm tracking

Worm tracking was performed at room temperature using a WormTracker 2.0 imaging system (Yemini *et al*. 2013). Immediately before the experiment, NGM plates were seeded with a thin lawn (10 μL) of OP50, on which five young adult animals at a time were transferred and recorded for 5 min. Tracking videos were analyzed using the WormLab software (MBF Bioscience).

### Defecation assay

Defecation assays were performed as described (Thomas 1990). Well-fed, young adult worms were singled to NGM plates with a thin lawn of OP50. Before starting the assay, worms were left to acclimate for 10 min. Plates were mounted on a Nikon Eclipse E400 microscope and worms were observed using the 20X DIC objective. A posterior body muscle contraction (pBOC) of the worm signaled the start of the observation period. For 10 min, the pBOC and enteric muscle contraction (EMC) events were counted, including the first pBOC at the start of the assay.

### Staining for lipids

Staining for lipids using Nile Red (NR) was performed following (Escorcia *et al*. 2018). L4 worms raised at 20 °C were harvested by washing with 1X PBST. Worms were pelleted by centrifugation at 560 x *g* for 1 min and supernatant discarded. To the pellet, 100 uL of 40% isopropanol was added and left to incubate for 3 min at room temperature. Worms were again centrifuged at 560 x *g* for 1 min and supernatant was carefully removed.

To prepare the NR stain, 6 uL of 5 mg/mL NR in 100% acetone was freshly added to 1 mL of 40% acetone. Working in the dark, 600 uL of NR stain was added and thoroughly mixed to each worm pellet by inverting the tubes. Samples were rotated in the dark for 2 h at room temperature. Worms were then pelleted at 560 x *g* for 1 min and the supernatant was removed. To remove excess NR stain, samples were incubated with 600 uL of PBST for 30 min in the dark. Samples were centrifuged at 560 x *g* for 1 min and all but 50 uL of supernatant was discarded. Worms were resuspended in the residual supernatant and imaged immediately after staining. Intensity of the lipid stain was measured in Fiji (Schindelin *et al*. 2012).

### Microscopy

Worms were immobilized using 100 mM sodium azide (NaN_3_) and mounted on 5% agarose pads on glass slides. Unless otherwise indicated, worms were imaged at the L4 or young adult stage. Images were acquired using either Zeiss LSM 980 confocal microscope or Axio Imager Z2 compound microscope at 40X magnification unless otherwise specified. Processing of images was done using the Zen (Zeiss) or Fiji (Schindelin *et al*. 2012) software. Fluorescence intensities of reporters were quantified using Fiji.

### Microfluidic locomotory analysis

The time-lapse images (1 frame/second, 5 h recording) for behavioral assays were acquired using a high-resolution, multi-well imaging system (Ji *et al*. 2024). We added 60 μL of liquid NGM buffer (same components as NGM but without agar or peptone) to each well of a 96-well plate (Corning, Inc.), then picked one Day 1 adult animal to each well. Where indicated, the NGM was supplemented with food *E. coli* OP50, and the bacteria was added to NGM to a final OD_600_ ≈ 1. Multiple genotypes or conditions being compared were assayed on the same plate.

Image data processing and analysis were performed as in a previous study (Churgin *et al*. 2017). Consecutive images were subtracted to generate difference images. We normalized the difference image with the average pixel intensity of the two subtracted images. To reduce noise, we applied a Gaussian filter with SD equal to one pixel to the difference image. We then applied a binary threshold of 0.6 to the filtered difference image to determine whether movement occurred at each pixel. We summed up all pixels of the binarized difference image and used the resulting value to define the activity. To calculate the dwelling fraction of worms, we time-averaged the activity data with a smoothing kernel of 5 s and generated a histogram of the time-averaged activity for each worm. We then employed a nonlinear least-squares fitting algorithm to fit each individual worm’s activity histogram to two exponential terms (2 free parameters each) and a Gaussian term (3 free parameters). The zero bin on the activity histogram (corresponding to quiescence) was excluded from the histogram before the fitting process. The dwelling fraction of each worm was calculated as the area under the two exponential curve components of the fit.

### Compensatory curvature response (CCR) assay

We employed a straight channel microfluidic device to perform the CCR behavioral assay as described in a previous study (Ji *et al*. 2023). The microfluidic device consists of 2000-μm-wide open areas and two parallel straight channels (60 μm width, 200 μm length, 80 μm height). During the assay, we transferred day-1 adults to a food-free NGM buffer for approximately 5 minutes to remove bacteria, and then pipetted the animals from the washing buffer into a microfluidic chamber filled with NGM buffer containing 0.1% (by mass) bovine serum albumin (for preventing worms from adhering to chamber surfaces). We recorded behavioral videos (30 frames/second) of each animal in the chamber for about 3 minutes performing free locomotion in open areas or constrained locomotion with their mid-body in the narrow channels.

The behavioral data from microfluidic experiments were analyzed using methods described previously (Ji *et al*. 2023). For each worm, the normalized bending curvature *K* is calculated as the product of body curvature *k* and the worm body length *L*. To quantify the effect of straight-channel constraint on worm bending curvature amplitude, the whole-body curvature amplitude during constrained locomotion was computed and compared with that of free locomotion. Specifically, we analyzed worm bending curvature dynamics during free locomotion to generate an averaged curvature amplitude profile, *A_free_*(*s*), as a function of body coordinate *s* defined such that s=0 at the head and s=1 at the tail. We divided video sequences of constrained movement into individual short sequences using a 3 s time window to analyze constrained locomotion. We manually marked the channel position in each image sequence to record the relative position of the constraint to the worm body. The normalized curvature change in response to mid-body constraint was calculated by considering only periods during which the anterior and posterior limits of the narrow channel were consistently within a body coordinate interval between 0.35 and 0.65. The resulting curvature dynamics were denoted as *K_const_*, and the maximum value of |*K_const_*(*s*, *t*)| in the time dimension was defined as the curvature amplitude profile of individual periods, denoted as 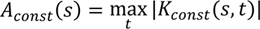. The normalized curvature change of each period was represented by *A_const_*(*s*)/*A_free_*(*s*) − 1. The normalized anterior curvature changes of individual periods was defined as 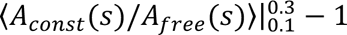 where 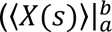 denotes the average of *X* over the interval [a, b]).

### Lifespan assay

Lifespan assays were performed as previously described (Amrit *et al*. 2014). One hundred (100) unhatched embryos were put on a plate, with 3 plates for each genotype. Throughout the experiment, plates were kept in dark boxes at 20 °C as continuous exposure to light has been shown to decrease worm lifespan (De MAGALHAES FILHO *et al*. 2018). Embryos were left to develop for three days until Day 1 of adulthood, after which 15 young adults were gently transferred to each plate, with seven plates for each genotype. The number of worms that were alive, dead, and censored were recorded every alternate day of adulthood. Censored worms are worms that die due to bagging, desiccation, contamination, or human error. Worms that are immobile and fail to respond to touch with a platinum pick are considered dead. Dead and censored worms were burned once they were recorded. For the first six days of adulthood, worms were transferred to a new plate every day to avoid contamination with their progeny. Once worms stopped producing progeny, usually after day 6 of adulthood, they were transferred to new plates every other day. Lifespan was recorded until death of the last worm, which usually occurred between 30-45 days of adulthood. In cases of contamination with mold or unwanted bacteria during the assay, all live worms were transferred to clean plates. Statistical analysis, Kaplan-Meier survival analysis, and Mantel-Cox log-rank test were performed using OASIS 2 software (https://sbi.postech.ac.kr/oasis2) (Han *et al*. 2016).

## RESULTS

### Group A bHLH family genes in the nervous system of *C. elegans*

The *C. elegans* genome encodes members of all six groups of bHLH genes (Groups A to F)(Ruvkun and Hobert 1998; Ledent *et al*. 2002; Simionato *et al*. 2008; Grove *et al*. 2009; Gyoja 2014). We focus here on members of group A since its striking expansion in metazoans (Gyoja 2014) provides a clear indication of their role in cell type diversification. There are 19 group A genes in *C. elegans*, including paralogues of the phylogenetically deeply conserved Ato, ASC, NeuroD, Neurogenin and MyoD proteins (**Fig.1A**). While several of these group A genes have been previously analyzed for their function within and outside the nervous system (Hallam *et al*. 2000; Portman and Emmons 2000; Fukushige and Krause 2005; Masoudi *et al*. 2018; Aquino-NUNEZ *et al*. 2020; Christensen *et al*. 2020; Filippopoulou *et al*. 2021; Masoudi *et al*. 2021), others have remained uncharacterized.

Examination of group A bHLH transcript presence in the whole nervous system single cell RNA atlas of *C. elegans*, established by the CeNGEN consortium (Taylor *et al*. 2021), reveals that only a subset of group A proteins are expressed in the mature *C. elegans* nervous system, and each in a very sparse manner in different mature neuron types (**Fig.1A**). These include several genes whose function in the nervous system differentiation have not previously been analyzed. We chose the five most strongly expressed and previously little studied genes from this category (**Fig.1B**), the *PTF1a/NATO3* ortholog *hlh-13*, the *NHLH1/2* ortholog *hlh-15* and the three bHLHb4/5 orthologs *hlh-17, hlh-31* and *hlh-32,* for an in-depth analysis of their expression and function. First, we used CRISPR/Cas9 genome engineering to tag each of these five genes with *gfp* to analyze the expression of the protein product of these loci throughout the whole animal during all life stages, thereby significantly expanding previous transcript and reporter transgene analysis. Second, we used existing or generated new deletion alleles, again using CRISPR/Cas9 genome engineering, to rigorously probe their function. Reporter and mutant alleles for each examined locus are schematically shown in **Fig.1C**.

### The bHLHb4 and bHLHb5 orthologs HLH-17 and HLH-32 proteins are expressed in neurons, but not glia

The *C. elegans* genome codes for three members of the Olig superfamily of transcription factors, *hlh-17, hlh-31* and *hlh-32*. In vertebrates, this family is defined by two closely related subtypes of genes, the highly interrelated, paralogous Olig1, Olig2 and Olig3 genes and the two highly related bHLHb4 (now also called bHLHe23) and bHLHb5 (now also called bHLHe22) gene paralogs. While the three *C. elegans* genes *hlh-17, hlh-31* and *hlh-32* have previously been considered as orthologs of the Olig subbranch of bHLH genes (Yoshimura *et al*. 2008), detailed sequence analysis, as well as orthology assignment tools demonstrate that *hlh-17, hlh-31* and *hlh-32* are more closely related to the bHLHb4/5 subbranch than to the Olig subbranch (Hu *et al*. 2011; Wang *et al*. 2017; Kim *et al*. 2018)(**Fig.1B, S1A-C**). Conversely, another *C. elegans* bHLH gene, *hlh-16,* appears to be more closely related to the Olig genes than *hlh-17, hlh-31* and *hlh-32* (**Fig.S1A-C**).

All three bHLHb4/5 paralogs cluster within a small interval (<150kb) on chromosome IV and *hlh-17* and *hlh-31* are direct neighbors (**Fig.S1D**). The *hlh-17* and *hlh-32* genes display identical bHLH domain-encoding sequences, indicating that they have the same protein dimerization and DNA binding properties (**Fig.S1B**). All three genes are the result of species-specific gene duplications within the nematode phylum. Reciprocal BLAST searches show that *C. remanei* has at least six members of the bHLHb4/b5 subfamily while *C. briggsae* has only a single representative (*Cbr-hlh-17*). Nematodes from other clades also do not harbor *hlh-17/31/32* duplications, leading us to conclude that an ancestral single bHLHb4/5 gene has undergone recent duplications within select members of the *Caenorhabditis* genus and then subsequently duplicated once in the vertebrate lineage, to generate bHLHb4 and bHLHb5. The related Olig-like gene *hlh-16* has not undergone gene duplications in nematode genomes, but it has been lost in *Drosophila*, which contains instead a single bHLHb4/b5 representative, somewhat misleadingly called Oli (Oyallon *et al*. 2012)(**Fig.1B, S1**).

The fusion of 5’ promoter sequences of the *hlh-17, hlh-31* and *hlh-32* genes to *gfp* reporters had shown that the *hlh-17* reporter construct is expressed in the CEP sheath (CEPsh) glia cells and unidentified motor neurons in the ventral nerve cord, while the *hlh-32* reporter fusion was expressed in an unidentified pair of head neurons; no expression could be discerned for the *hlh-31* promoter fusion construct (Mcmiller and Johnson 2005; Yoshimura *et al*. 2008). Promoter fusion constructs can miss *cis-* regulatory elements and, particularly in the case of the *hlh-32/17/31* locus, the genomic rearrangements leading to the duplication of the ancestral gene may have resulted in unusual arrangements of *cis-*regulatory elements. To eliminate these concerns, we tagged each of the three endogenous loci with *gfp*, using CRISPR/Cas9 genome engineering. The *gfp* tagged loci show expression patterns that are indeed different from the previously reported promoter fusions. We find that HLH-17::GFP and HLH-32::GFP protein are exclusively detectable in three neuron classes, the bilaterally symmetric glutamatergic AUA interneuron class and two B-class cholinergic ventral cord motor neurons, DB2 and VB2 (**Fig.2**). Together with a few other head and neck muscle-innervating motor neurons, these motor neurons are part of the retrovesicular ganglion (RVG), which is located at the anterior end of the ventral nerve cord. DB2 and VB2 are the most anteriorly located B-type motor neurons in the RVG and have lineage histories that are distinct from other DB and VB class members (Sulston and Horvitz 1977; Sulston *et al*. 1983). Based on scRNAseq data (Taylor *et al*. 2021; Smith *et al*. 2024), DB2 and VB2 are also molecularly subtly distinct from other DB and VB class members. Due to their location, the RVG motor neurons are analogous to the branchial motor neurons in the brain stem of vertebrates, but little is known about how RVG motor neuron subtypes are made to become different from other MNs of the same type.

**Fig. 2:**
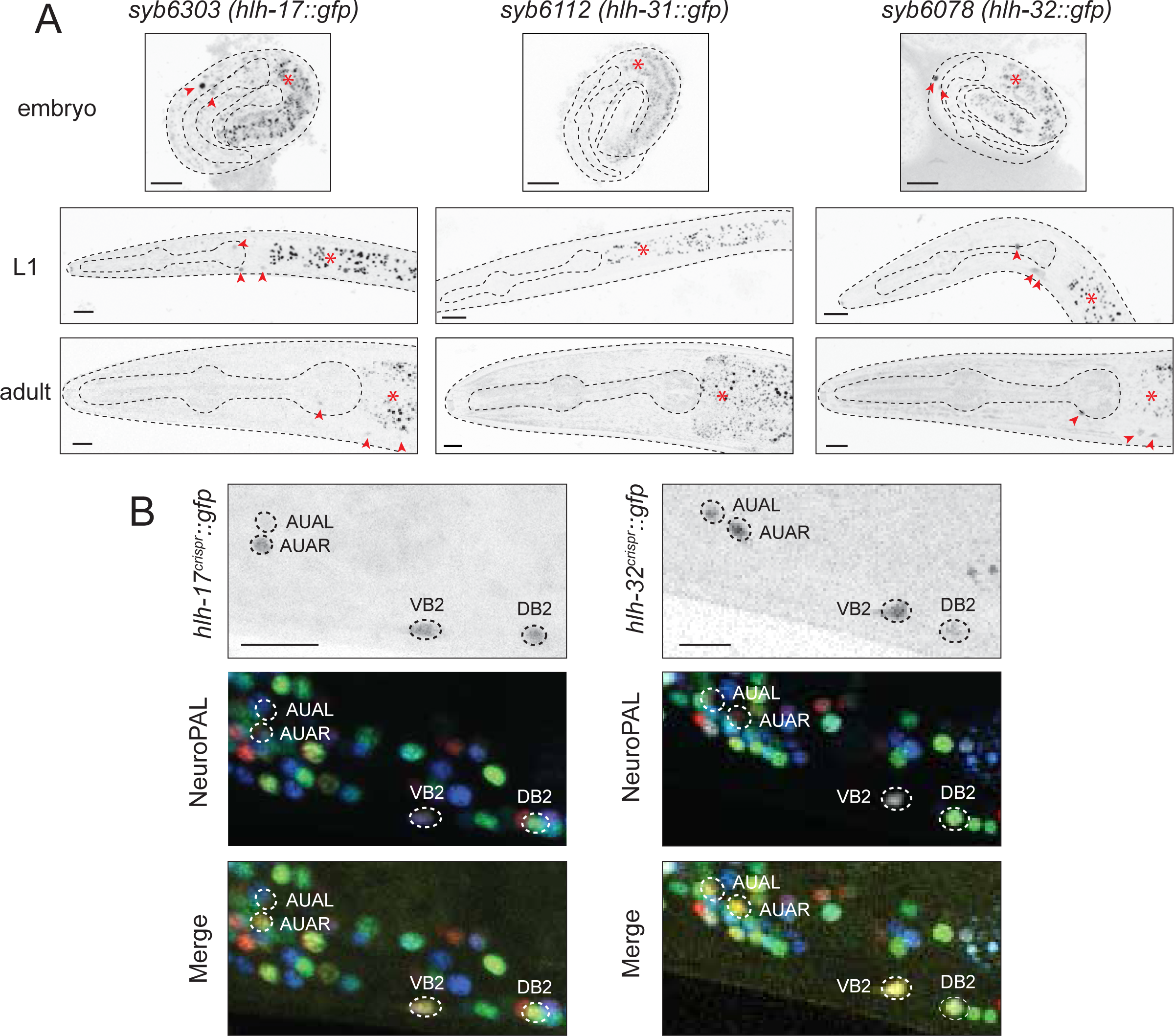
Expression of *hlh-17, hlh-31* and *hlh-32* reporter alleles. **A:** Expression of CRISPR/Cas9-engineered reporter alleles, *hlh-17(syb6303)*, *hlh-31(syb6112)*, and *hlh-32(syb6078)* over the course of development (see Fig.1C for reporter allele). HLH-17::GFP and HLH-32::GFP can be detected in head neurons (red arrowheads), while no HLH-31::GFP signal could be detected anywhere at any stage. Asterisks indicate intestinal autofluorescence. Representative images for reporter alleles are shown with 10 μm scale bar. **B:** Overlay with the NeuroPAL *(otIs669)* transgene demonstrates *hlh-17(syb6303)* and *hlh-32(syb6078)* expression in the AUA, VB2 and DB2 neurons. Representative close-up views for reporter alleles are shown with 10 μm scale bar.

Expression of HLH-17::GFP and HLH-32::GFP in AUA, DB2 and VB2 is observed throughout all larval and adult stages, commences when those neurons are generated (**Fig.2**) and is consistent with scRNAseq data (Taylor *et al*. 2021) (**Fig.1A**). Notably, however, we detected no HLH-17::GFP protein expression in CEPsh glia (or any other glia) at any developmental stage, indicating that *hlh-17* transcripts, detected by scRNAseq analysis (Taylor *et al*. 2021) and inferred from transcriptional reporter constructs (Mcmiller and Johnson 2005; Yoshimura *et al*. 2008) may not become translated into detectable protein levels in the CEPsh glia. We have occasionally observed similar transcript and protein discordance for homeobox genes (Reilly *et al*. 2020; Taylor *et al*. 2021).

In contrast to HLH-32 and HLH-17, expression of HLH-31::GFP, encoded by the gene that directly neighbors HLH-17 (and which displays a somewhat degenerate bHLH domain sequence; **Fig.S1**), was not detectable in our analysis at any stage in any cell within or outside the nervous system. scRNA transcriptome analysis detects transcripts for *hlh-31* in the DB2 and VB2 neurons (like its neighboring *hlh-17* gene), and also in the CAN neurons (Taylor *et al*. 2021) (**Fig.1A**). However, as with *hlh-17* transcripts in the CEPsh cells (also detected by scRNAseq), no readily detectable protein appears to be made from these *hlh-31* transcripts.

### Removal of *hlh-17, hlh-31* and *hlh-32* does not obviously affect glia differentiation

To assess potential functions of *hlh-17, hlh-31* and *hlh-32,* we generated single, double and triple mutant strains using CRISPR/Cas9 genome engineering. Unlike the previously generated *hlh-17, hlh-31* and *hlh-32* mutant alleles, which left parts of each gene intact (Yoshimura *et al*. 2008), our engineered alleles each deleted the entire respective locus (**Fig.1C**). Due to overlaps in gene expression, and the sequence identity of the bHLH domains (**Fig.S1B**), we considered possible functional redundancies and therefore started our analysis by examining *hlh-17/hlh-31/hlh-32* triple null mutant animals. These were generated in two consecutive genome editing rounds, first removing the two adjacent *hlh-17* and *hlh-31* loci (*syb6635)* and then removing the more distal *hlh-32* locus (*syb7713)* in the background of the *syb6635* allele. From here on we refer to these *hlh-32(syb7773)hlh-17hlh-31(syb6635)* triple null mutants as “*hlh-17/31/32^null^”*. These triple null mutant animals are viable, fertile and display no obvious morphological or behavioral abnormalities.

Consistent with the absence of HLH-17/31/32 protein expression in the CEPsh glia, we observed no effects of *hlh-17/31/32^null^* mutants on CEPsh glia differentiation, as assessed by examination of CEPsh morphology and marker gene expression (**Fig.S2**).

### *hlh-17* and *hlh-32* are required for AUA neuron differentiation

To examine AUA neuron identity specification, we used the NeuroPAL transgene, in which this neuron is marked with two distinct fate markers, *eat-4/VGLUT* and *mbr-1* (Yemini *et al*. 2021). We find that the NeuroPAL color code of the AUA neuron pair is not obviously changed in *hlh-17/31/32^null^*animals (**Fig.3A**). Examining other AUA markers, we first investigated expression of the neuropeptide-encoding *flp-8* and *flp-32* genes (Kim and Li 2004; Taylor *et al*. 2021). We found that *flp-8*, but not *flp-32*, expression is affected in triple null mutants (**Fig.3B,C**). Neither *hlh-32(ot1347)* single mutants, nor *hlh-17hlh-31(syb6635)* double mutant recapitulated the *flp-8* expression defects (**Fig.3C**), indicating that AUA-expressed *hlh-32* and *hlh-17* indeed act redundantly to specify *flp-8* expression in AUA. This is not surprising given that both proteins are co-expressed and display an identical amino acid sequence of their bHLH domain.

**Fig. 3:**
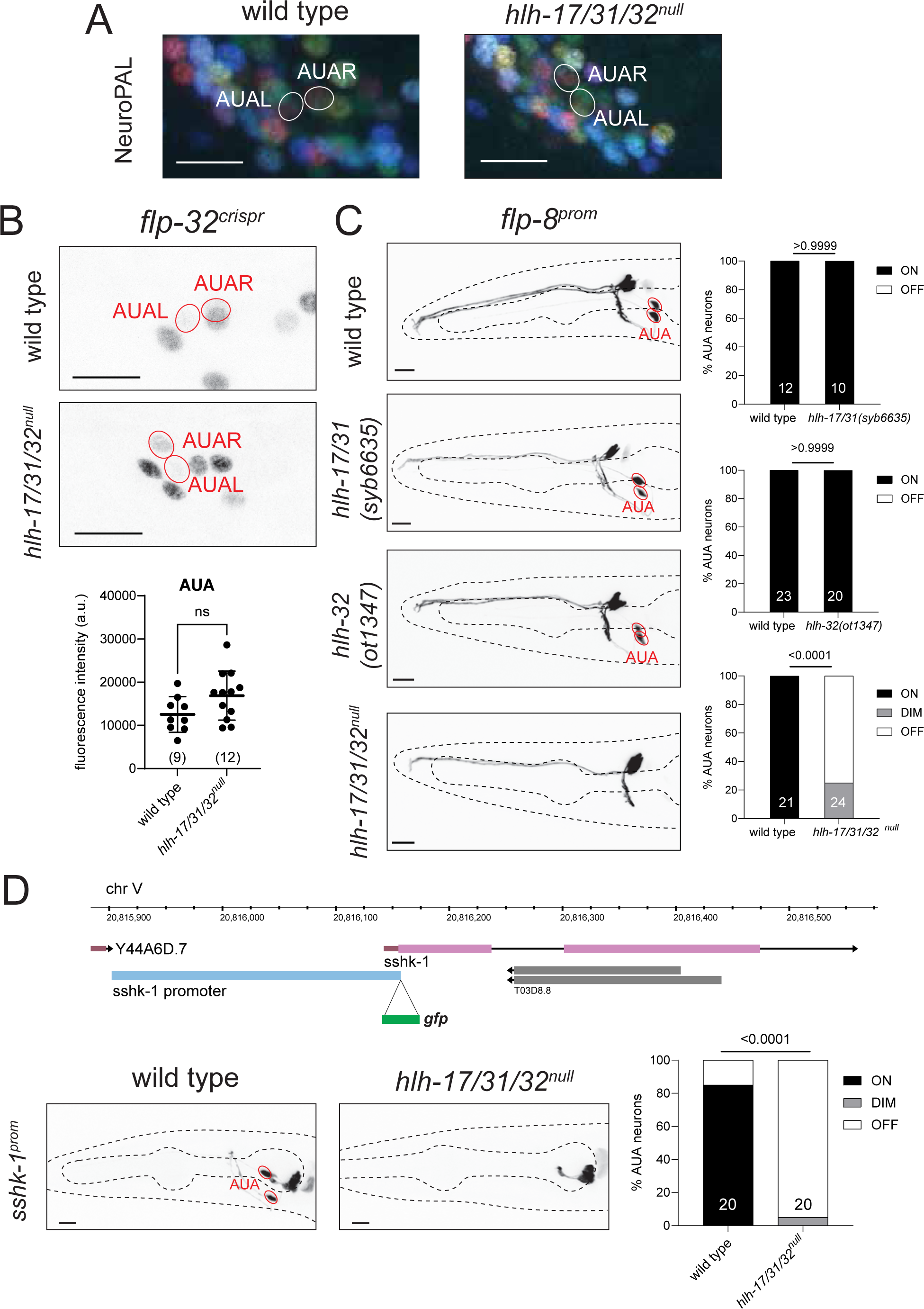
AUA cell fate analysis of *hlh-17, hlh-31* and *hlh-32* mutant animals. **A,B:** Triple *hlh-17/31/32*^null^ mutants do not show a change of NeuroPAL *(otIs669)* coloring of AUA (panel A) or expression of *flp-32(syb4374)* in AUA (panel B). Shown are representative images of wild type and mutant animals with 10 μm scale bar. Statistical analysis was performed using unpaired t-test. Individual points represent averages of the fluorescence intensities of the AUA neuron pair within an individual. The number of animals scored are enclosed in parentheses. Error bars indicate standard deviation of the mean. ns: not significant. **C:** Expression of *flp-8* reporter *(ynIs78)* in AUA is affected only in *hlh-17/31/32*^null^ triple mutants. Individual *hlh-17/31(syb6635)* and *hlh-32(ot1347)* mutants do not show loss of the *flp-8* reporter. Shown are representative images of wild type and mutant animals with 10 μm scale bar. Number of animals scored are shown within each bar. P-values were calculated using Fisher’s exact test. **D:** A novel fate reporter for AUA was generated by fusing the promoter directly upstream *sshk-1* with *gfp (otEx8199)*. *hlh-17/31/32*^null^ triple mutants lose expression of *sshk-1* reporter in AUA. Representative images of wild type and mutant animals are shown with 10 μm scale bar. Number of animals scored are shown within each bar. P-values were calculated using Fisher’s exact test.

We generated another identity marker for the AUA neuron by interrogating the CeNGEN scRNA atlas for transcripts with selective, strong expression in AUA. We considered the *T03D8.7* locus, which codes for a small secreted protein with a ShKT domain that, in other species, acts as a toxin against pathogens by inhibiting potassium channels (Tudor *et al*. 1996). We named this gene *sshk-1* (for small ShKT domain) and found that a transcriptional reporter that encompasses the entire intergenic region to its preceding gene is indeed expressed exclusively in AUA within the entire nervous system; outside the nervous system expression is also observed in the exocrine pharyngeal gland cells, consistent with this protein having a potentially protective function against pathogens (**Fig.3D**). We found that expression of the *sshk-1^prom^* reporter transgene is strongly affected in the AUA (but not gland cells) of *hlh-17/31/32^null^*mutant animals (**Fig.3D**). In those animals where residual *sshk-1^prom^* expression was visible, AUA neurite morphology appeared normal.

### *hlh-17* and *hlh-32* diversify VB1/VB2 motor neuron subtype identity

We interrogated motor neuron identity specification using again *hlh-17/31/32^null^* triple mutant animals because of (a) the redundant functions of co-expressed *hlh-32* and *hlh-17* we had revealed above in the AUA fate analysis and (b) because of the theoretical possibility that even though *hlh-31* showed no expression in wild-type animals, it may be upregulated in *hlh-32; hlh-17* double mutants. Between the motor neurons that express *hlh-17* and *hlh-32*, VB2 and DB2, we focused our analysis on VB2 because more identity markers are available for VB2 than for DB2.

We first examined the effect of the *hlh-17/31/32^null^* mutation on NeuroPAL, in which defects in neuronal specification can be detected through changes in neuron coloring conferred by specific sets of promoters (Yemini *et al*. 2021). We observed that in *hlh-17/31/32^null^*animals, VB2 adopts NeuroPAL coloring that resembles another VB class member, VB1 (**Fig.4A**). The change in NeuroPAL coloring of VB2 can be attributed to loss of *acr-5* expression, which normally distinguishes VB2 from VB1 (Yemini *et al*. 2021). Previous cell fate marker analysis from our lab (Kerk *et al*. 2017; Sun and Hobert 2021; Yemini *et al*. 2021), as well as recent scRNAseq analysis (Taylor *et al*. 2021; Smith *et al*. 2024) revealed a number of additional genes that are differentially expressed in VB2 versus VB1, which allowed us to test whether *hlh-17/31/32* may indeed work to distinguish the identity of these neurons. We first interrogated markers shared by VB2 and VB1 but expressed at different levels, according to scRNA transcriptomes (Taylor *et al*. 2021; Smith *et al*. 2024), namely the neuropeptides *flp-32* and *flp-7.* Using CRISPR/Cas9-engineered reporter alleles, we found that the difference in expression level of these markers that exists between VB2 and VB1 in wild type animals is abolished in *hlh-17/31/32^null^*mutants (**Fig.4B-C**).

**Fig. 4:**
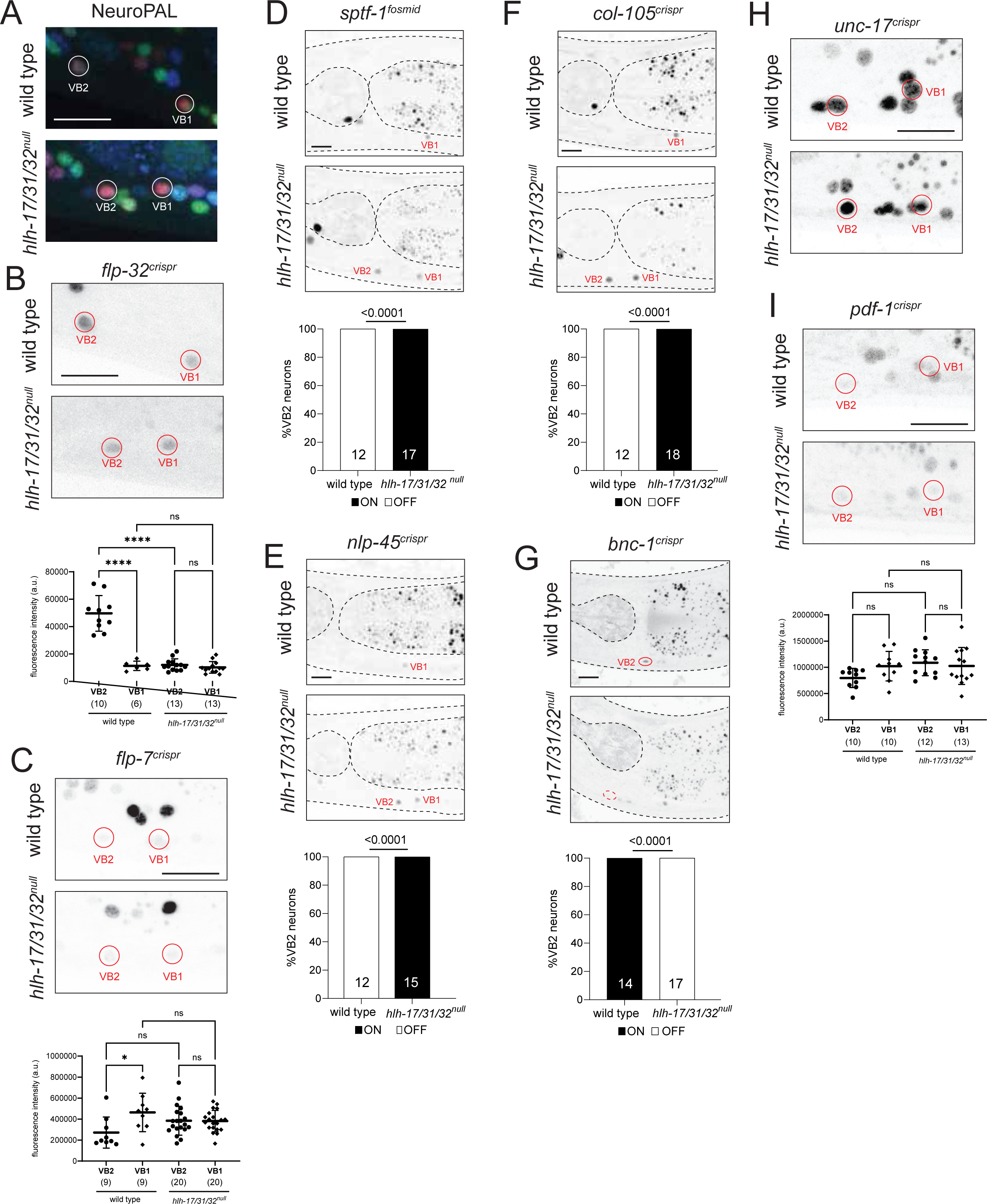
VB2 cell fate analysis of *hlh-17, hlh-31* and *hlh-32* mutant animals. **A:** In *hlh-17/31/32*^null^ mutants, VB2 adopts a VB1-like NeuroPAL *(otIs669)* coloring. **B:** In *hlh-17/31/32*^null^ mutants, *flp-32(syb4374)* reporter allele expression in VB2 is decreased, reaching the same level of expression as that in VB1. **C:** The difference in *flp-7(syb5413)* reporter allele expression levels between VB2 and VB1 is abolished in *hlh-17/31/32*^null^ mutants. **D:** Expression of the *wgIs707* fosmid-based reporter for transcription factor *sptf-1* is normally restricted to VB1 but becomes derepressed in VB2 in *hlh17/31/32*^null^ animals. **E:** The *nlp-45(ot1032)* reporter allele marks VB1 alone in wild type but is ectopically expressed in VB2 in *hlh-17/31/32*^null^ animals. **F:** The *col-105(syb6767)* reporter allele is a marker for VB1 that is ectopically expressed in VB2 in *hlh-17/31/32*^null^ animals. **G:** The reporter allele for transcription factor *bnc-1(ot845)* loses expression in VB2 in *hlh-17/31/32*^null^ animals. **H:** *hlh-17/31/32*^null^ mutants retain strong expression of *unc-17(syb4491)* in VB2 and VB1. **I:** Expression of the *pdf-1(syb3330)* reporter allele in VB2 and VB1 remain unchanged in *hlh-17/31/32*^null^ mutants. Representative images of wild type and mutant animals are shown with 10 μm scale bar. For panels B, C, and E, statistical analysis was performed using one-way ANOVA with Tukey multiple comparisons test. Each point represents the corresponding neuron from one individual and the number of animals scored are enclosed in parentheses. Error bars indicate standard deviation of the mean. ****p ≤ 0.0001, *p ≤ 0.05, ns: not significant. For panels F-I, statistical analysis was performed using Fisher’s exact test, with the number of animals scored shown within each bar.

If VB2 indeed acquires VB1 fate, markers exclusive to VB1 should become ectopically expressed in the VB2 neurons of *hlh-17/31/32^null^*mutants. To examine such a potential VB2-to-VB1 transformation, we sought markers normally present only in VB1 but not VB2. A fosmid-based reporter transgene of the *sptf-1* Zn finger transcription factor had previously been shown to be expressed in VB1, but not VB2 (Taylor *et al*. 2021) **(Fig.4D)**. Moreover, we found that a *nlp-42* neuropeptide reporter allele, previously described to be in either VB1 or VB2 (Sun and Hobert 2021), is expressed in VB1, not VB2 (**Fig.4E**). In addition, based on scRNAseq data (Taylor *et al*. 2021; Smith *et al*. 2024), we CRISPR/Cas9-engineered a reporter allele for the collagen *col-105,* and confirmed expression in VB1, not VB2 (**Fig.4F**). Armed with these three VB1(+) VB2(-) markers, we indeed found that all three (*sptf-1, nlp-45, col-105*) become ectopically expressed in the VB2 neuron of *hlh-17/31/32^null^*triple mutant animals (**Fig.4D-F**). Lastly, a Zn finger transcription factor, *bnc-1,* normally expressed in VB2 (and all other B-type motor neurons), but not VB1 (Kerk *et al*. 2017), fails to be properly expressed in VB2 in *hlh-17/31/32^null^* mutants (**Fig.4G**).

Markers that are expressed at similar levels between VB2 and VB1, such as reporter alleles for the pan-cholinergic marker *unc-17/VAChT* or the *pdf-1* neuropeptide, are unaffected in *hlh-17/31/32^null^*mutants (**Fig.4H-I**). Together with the observation that the pan-B-type motor neuron marker *acr-2* within NeuroPAL remains expressed in *hlh-17/31/32^null^* mutants (**Fig.4A)**, our results suggest that *hlh-17/31/32* are neither required for the generation of VB2 nor for the adoption of B-motor neuron fate but may control the diversification of VB2 away from VB1 identity.

We conclude that VB2-expressed *hlh-17/31/32* ensure the diversification of VB2 and VB1 identity by promoting expression of VB2 features and repressing VB1-specific features, without affecting overall B-type motor neuron identity specification. To our knowledge this is the first description of a molecular mechanism of B-type motor neuron diversification within the retrovesicular ganglion.

### Behavioral analysis of *hlh-17 hlh-31 hlh-32* triple mutant animals

*hlh-17* mutants were previously reported to display defects in the defecation motor program, which was ascribed to *hlh-17* function in the CEPsh glia (Bowles and Johnson 2021). We used our more unambiguous *hlh-17* null allele, in combination with the *hlh-31* and *hlh-32* null alleles, to avoid any potential issues of genetic redundancies among these genes, to confirm these defects. Testing both the posterior body wall contraction (pBOC) and the enteric muscle contraction (EMC) step of the defecation motor program, we observed no defects in *hlh-17/31/32^null^* triple mutant animals (**Fig.S3A**).

We also examined locomotory behavior of *hlh-17/31/32^null^* mutant animals using an automated Wormtracker system (Yemini *et al*. 2013) and observed a number of defects, including reversal behavior and speed (**Fig.S3B**). We have not pursued the question of whether those defects are due to AUA and/or VB2 differentiation defects. We note, however, that whole brain activity recordings have shown striking correlations of the activity patterns of AUA with the command interneuron AVA (Yemini *et al*. 2021), which controls several locomotory features that we find to be defective in *hlh-17/31/32^null^* mutant animals.

### Exclusive expression of the PTF1a ortholog HLH-13 in the octopaminergic RIC interneurons

The *C. elegans* genome encodes one ortholog of the two paralogous vertebrate PTF1a and NATO3 (aka FERD3L) bHLH genes, called *hlh-13* (Ledent *et al*. 2002; Simionato *et al*. 2008; Bao *et al*. 2017)(**Fig.1B**). A previous analysis of *C. elegans hlh-13/PTF1a* suggested expression in dopamine neurons, based on a small reporter transgene that contained only 2.1 kilobases upstream of the gene (Liachko *et al*. 2009; Bou DIB *et al*. 2014). However, the reported position of the GFP(+) cells in larval/adult animals is not fully consistent with expression in dopamine neurons and scRNAseq data also suggests expression in, at most, a subset of dopaminergic neurons, plus other neurons that showed no expression of the previous reporter transgene (TAYLOR *et al*.

2021). To resolve these issues, we tagged the endogenous *hlh-13/PTF1a* gene locus with *gfp,* using CRISPR/Cas9 engineering (**Fig.1C**). Using the NeuroPAL cell ID tool (Yemini *et al*. 2021), we observed strong HLH-13 protein expression exclusively in the sole octopaminergic neuron class in *C. elegans,* RIC (Alkema *et al*. 2005), throughout all larval and adult stages (**Fig.5A**). This is consistent with scRNAseq data of L4 stage animals (Taylor *et al*. 2021)(**Fig.1A**). RIC is also among the very few neuron classes that continuously express the E/Da protein HLH-2 throughout postembryonic life (**Fig.1A**)(Krause *et al*. 1997; Masoudi *et al*. 2022). Together with their documented capacity to interact in a yeast 2-hybrid assays (Grove *et al*. 2009), this co-expression indicates that like its vertebrate ortholog (Rose *et al*. 2001), HLH-13/PTF1A may heterodimerize with the E protein HLH-2 *in vivo*.

**Fig. 5:**
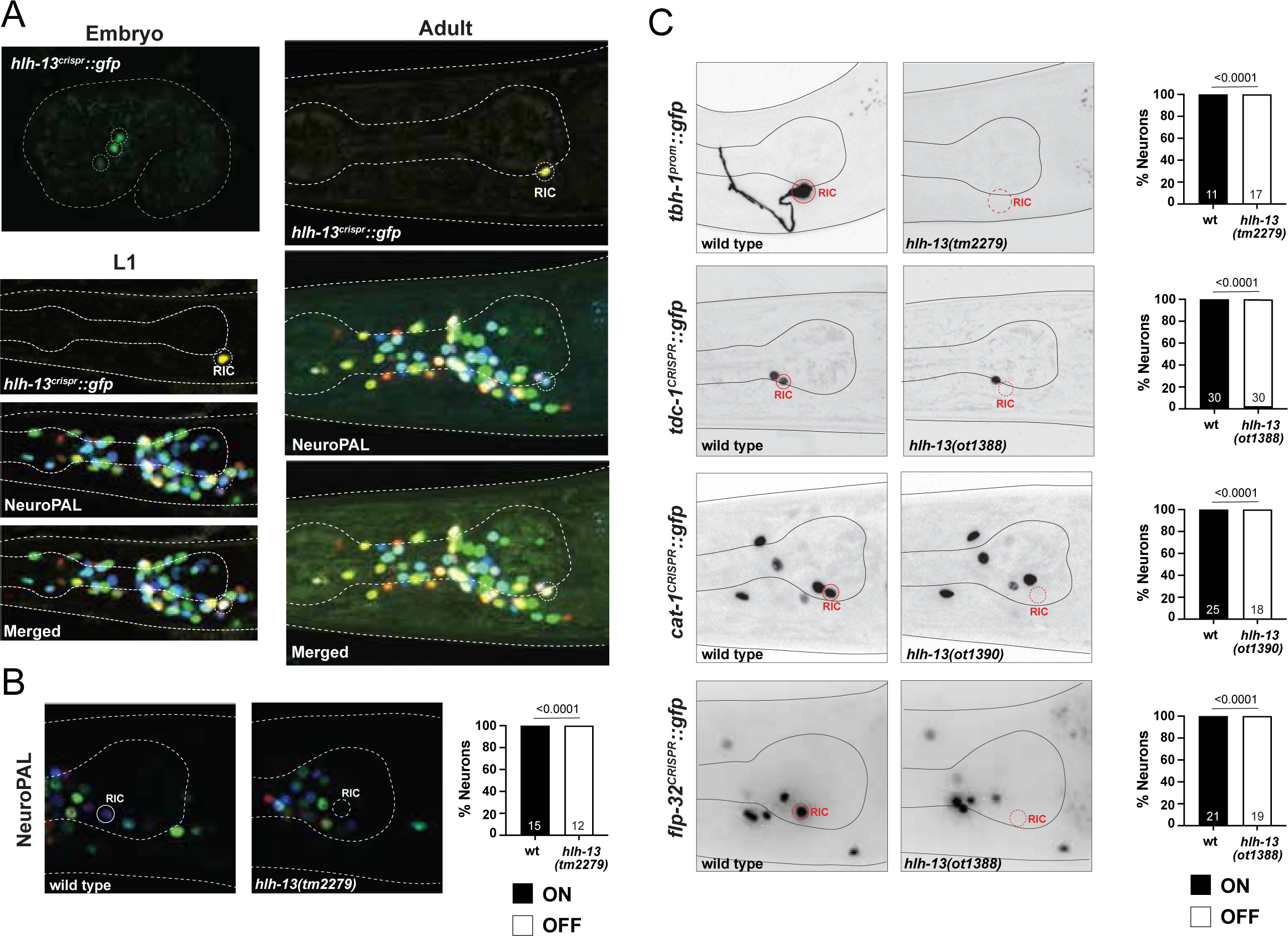
PTF1a and NATO3 ortholog HLH-13 is expressed in RIC and controls its differentiation. **A**: Expression of the *hlh-13* CRISPR/Cas-9-engineered reporter allele *syb7685* over the course of development. Overlay with the NeuroPAL transgene *otIs669* shows expression in RIC neurons. **B:** Representative pictures and quantification showing loss of NeuroPAL (*otIs669*) coloring in *hlh-13/PTF1a* mutants. Animals were scored at the young adult stage. Statistical analysis was performed using Fisher’s exact test. N is indicated within each bar and represents number of animals scored. **C:** Representative pictures and quantification showing loss of cell fate markers in *hlh-13/PTF1a* mutants. Reporter genes used are promoter fusion transgene for *tbh-1* (*nIs107*) and CRISPR/Cas9-engineered reporter alleles for *tdc-1*(*syb7768*), *cat-1*(*syb6486*) and *flp-32*(*syb4374*). Because *hlh-13* and *cat-1* are linked, the exact same mutation as *hlh-13(ot1388)* (Fig.1C) was generated in the background of the *cat-1(syb6486)* reporter; this allele is *hlh-13(ot1390).* Animals were scored at the young adult stage. Statistical analysis was performed using Fisher’s exact test. N is indicated within each bar and represents number of animals scored.

In the embryo we observe expression in 3 neuron pairs that resolves to expression in a single neuron pair (RIC) before hatching (**Fig.5A**). Since *hlh-13/PTF1a* scRNA transcripts are observed in embryonic (but not postembryonic) CEP neurons (Packer *et al*. 2019; Taylor *et al*. 2021), we crossed the *hlh-13/PTF1a* reporter allele with a transgenic line that expresses RFP in dopaminergic neurons including CEP, but a lack of overlap of the fluorescent signals ruled out that these neurons are the embryonic CEP neurons. Based on embryonic lineaging data of a *hlh-13/PTF1a* reporter transgene by the EPIC project (Murray *et al*. 2012), these HLH-13(+) cells may be the sister of the RIC neurons that are fated to die during embryogenesis and the two cousins of the RIC neurons, the SIBD neurons.

### Octopaminergic RIC neurons fail to properly differentiate in *hlh-13/PTF1a* mutant animals

We used two mutant alleles to analyze *hlh-13/PTF1a* function: (1) a deletion mutant, *tm2279,* generated by the National BioResource Project (NBRP) knockout consortium, which deletes most of the locus, including most of its bHLH domain and (2) *ot1388*, a complete locus deletion that we generated by CRISPR/Cas9 genome engineering (**Fig.1C**). Animals carrying either allele are viable, fertile and display no obvious morphological abnormalities. We assessed the function of *hlh-13/PTF1a* in neuronal differentiation by again using the NeuroPAL transgene. The RIC neuron class, the only cell type with strong and consistent postembryonic expression, display a loss of both markers on the NeuroPAL transgene, *ggr-3* and *mbr-1* (**Fig.5B**).

We extended our analysis of the RIC interneurons differentiation defects by examining additional markers of their octopaminergic identity, including the two biosynthetic enzymes TDC-1 and TBH-1 (Alkema *et al*. 2005). TDC-1, an aromatic amino acid decarboxylase is exclusively expressed in the octopaminergic RIC and tyraminergic RIM neurons to convert tyrosine to tyramine and TBH-1, a tyramine hydroxylase, is exclusively expressed in RIC to convert tyramine to octopamine (Alkema *et al*. 2005). We found that expression of both a *tbh-1* reporter transgenes and a *tdc-1* reporter allele (Wang *et al*. 2024) is eliminated in the RIC neurons of *hlh-13/PTF1a* mutants (**Fig.5C**). Moreover, a CRISPR/Cas9-engineered reporter allele of the monoaminergic vesicular transporter CAT-1/VMAT (Wang *et al*. 2024) also fails to be expressed in the RIC neuron of *hlh-13/PTF1a* (**Fig.5C**). Expression of *cat-1/VMAT* is unaffected in dopaminergic neurons in *hlh-13/PTF1a* mutants (**Fig.5C** and data not shown), consistent with *hlh-13/PTF1a* not being expressed in dopaminergic neuron development.

The RIC neuron pair also expresses a unique signature of neuropeptide-encoding genes (Taylor *et al*. 2021). We used a CRISRP/Cas9-engineered reporter allele of one of them, *flp-32* (Cros and Hobert 2022), and found that its expression is eliminated in *hlh-13/PTF1a* null mutants (**Fig.5C**).

We examined the fate of other neurons that are predicted to express much lower levels of *hlh-13/PTF1a* mRNA based on embryonic or postembryonic scRNAseq data, including all dopaminergic neurons, and found no differentiation defects using NeuroPAL. The above-mentioned *flp-32* RIC marker is also expressed in ALA (in which *hlh-13* transcript but no HLH-13::GFP protein can be detected), but we observed no *flp-32* expression defects in ALA. We conclude that *hlh-13/PTF1a* selectively affects RIC neuron differentiation.

### Physiological and behavioral consequences of *hlh-13/PTF1a* loss

The *hlh-13-*expressing RIC neuron is the only neuron that produces octopamine (Alkema *et al*. 2005), an endocrine monoaminergic regulator, analogous to vertebrate norepinephrine, that links nutrient cues to lipolysis to maintain energy homeostasis (Tao *et al*. 2016). We confirmed that loss of *tbh-1* results in an upregulation of lipid stores in the intestine, both under well-fed and starved conditions (**Fig.6A**), as previously reported (Tao *et al*. 2016). Consistent with *hlh-13/PTF1a* controlling RIC differentiation, and hence octopamine synthesis and release, we find that *hlh-13/PTF1a* mutant animals also display improper fat accumulation in the intestine, both under well-fed and starved conditions **(Fig.6A**).

**Fig. 6:**
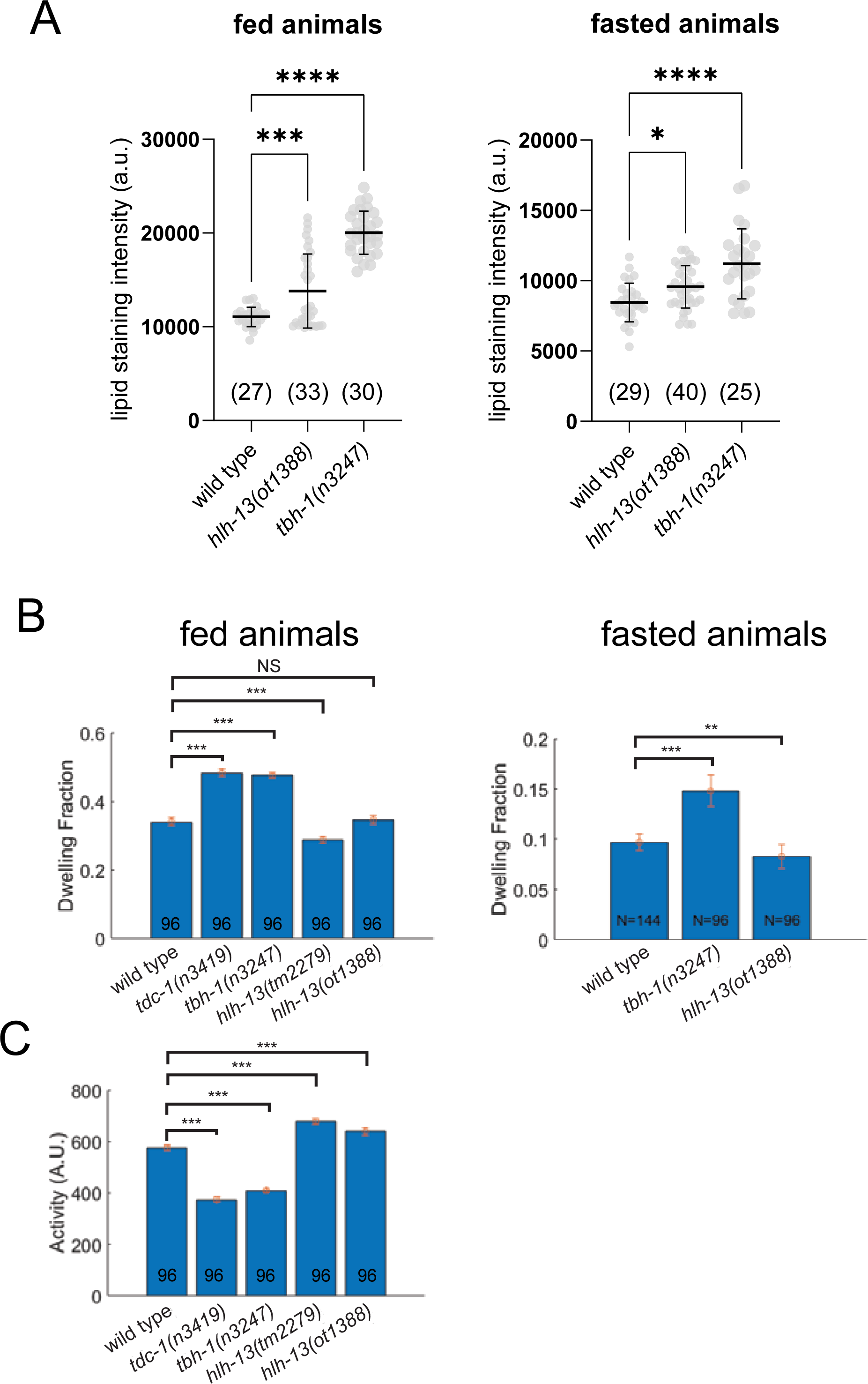
Physiological consequences of loss of the PTF1a and NATO3 ortholog *hlh-13*. **A:** *hlh-13(ot1388)* and *tbh-1(n3247)* mutant animals display comparable Nile Red lipid staining defects in well fed and in starved animals. Statistical analysis was performed using one-way ANOVA with Dunnett’s multiple comparisons test. Error bars indicate standard deviation of the mean. Number of animals scored for each genotype is enclosed in parentheses. ****p ≤ 0.0001, ***p ≤ 0.001, *p ≤ 0.05. **B,C:** Two independent *hlh-13/PTF1a* mutant alleles do not phenocopy the previously reported (Churgin *et al*. 2017)(and herein repeated) locomotory defects of *tbh-1(n3247)* mutant animals, indicating that the cellular source of octopamine required for proper locomotory behavior may not be the AVK neurons. Error bars indicate mean ± SEM. ***P < 0.001, **P<0.01, ns: not significant, compared with wild-type, Dunnett’s multiple comparison tests.

As another adaptation to changes in food availability, octopamine also controls dwelling behavior. Quantitative locomotory analysis of feeding *C. elegans* showed that *tbh-1* mutant animals exhibit a larger fraction of dwelling and smaller fraction of roaming compared to wild type worms (Churgin *et al*. 2017). Interestingly, this difference is not recapitulated in *hlh-13/PTF1a* mutant animals (**Fig.6B-C**), which exhibit roaming and dwelling at rates similar to that of wild type worms. Since octopamine is produced not only in the RIC neurons, but also in gonadal sheath cells (Alkema *et al*. 2005), we surmise that octopamine from the gonadal sheath cells may be sufficient for normal modulation of dwelling behavior.

### Exclusive expression of the NHLH1 and NHLH2 ortholog HLH-15 in two neuron classes

Previous phylogenetic analysis has shown that the two paralogous vertebrate bHLH genes NSCL1/NHLH1/HEN1 and NSCL2/NHLH2/HEN2 have a single *C. elegans* ortholog, *hlh-15* (Ledent *et al*. 2002; Simionato *et al*. 2008; Bao *et al*. 2017)(**Fig.1B**). A promoter-based transgene of *hlh-15/NHLH* was previously reported to be expressed in the tail DVA neuron and either the RIF or RIG head interneurons (Grove *et al*. 2009).

We used the CRISPR/Cas9 system to tag the *hlh-15/NHLH* locus with *gfp* (**Fig.1C**) and again used the NeuroPAL cell ID tool to assess the sites of *hlh-15::gfp* expression (Yemini *et al*. 2021). Throughout all postembryonic stages and adulthood, we observe HLH-15/NHLH protein expression exclusively in three neurons, the two peptidergic bilateral AVK interneurons AVKL and AVKR as well as the unpaired cholinergic tail interneuron DVA (**Fig.7A-C**). The exclusive expression in these two neuron classes commences at around the comma/1.5 fold stage and persists throughout all larval and adult stages. Both the embryonic and postembryonic reporter allele expression is consistent with scRNAseq data from embryos and L4 stage animals (Packer *et al*. 2019; Taylor *et al*. 2021)(**Fig.1A**). We re-examined the previously published promoter fusion construct and found that this construct is indeed also expressed in DVA and the AVK neurons (and not in RIF or RIG, as previously described; (Grove *et al*. 2009)).

**Fig. 7:**
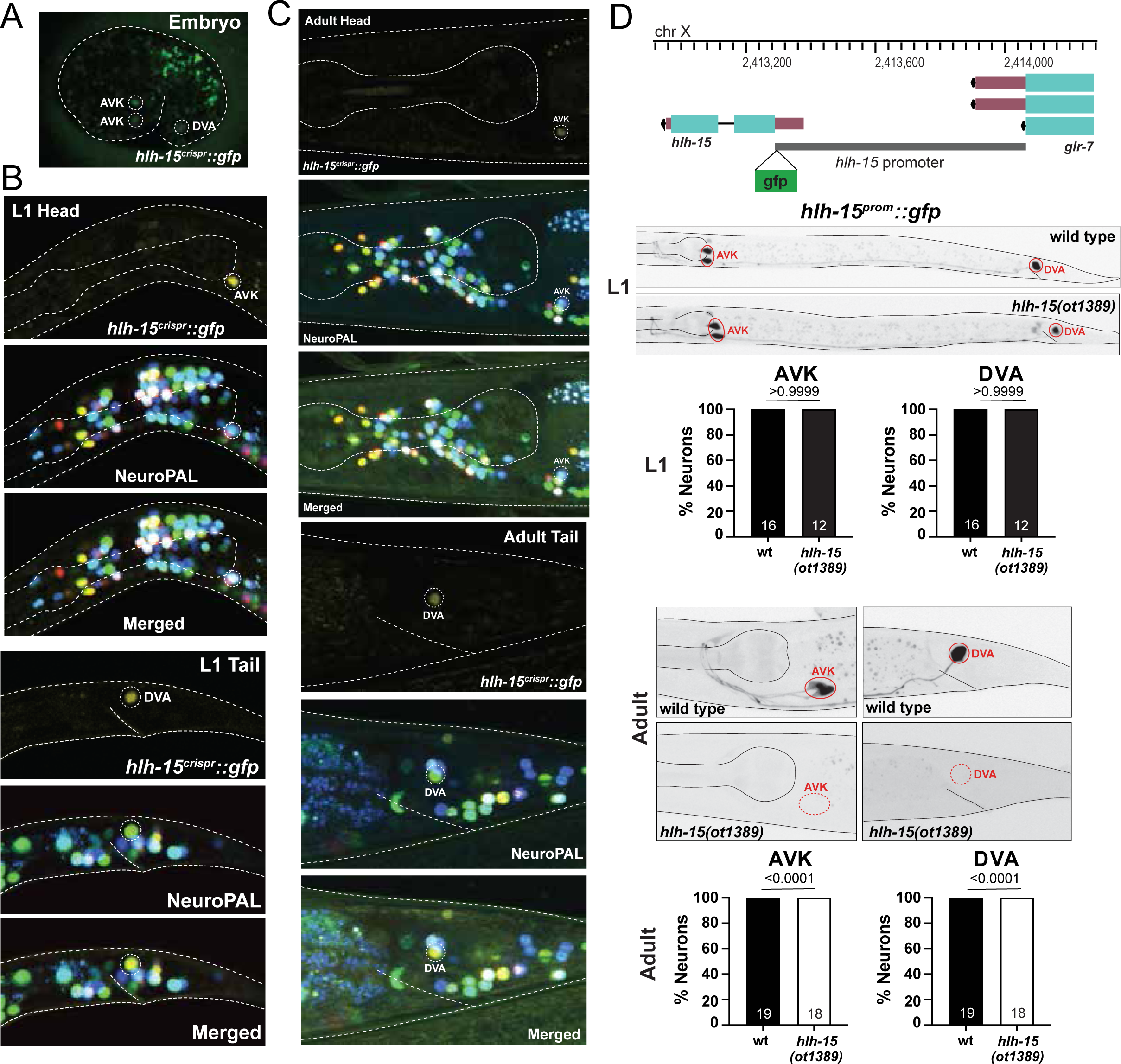
NHLH1 and NHLH2 ortholog HLH-15 is expressed in AVK and DVA neurons. **A-C:** Expression of the *hlh-15/NHLH* CRISPR/Cas-9-engineered reporter allele *syb7688* over the course of development. Overlay with the NeuroPAL transgene *otIs669* shows expression in AVK and DVA neurons. **D**: *hlh-15/NHLH* autoregulates its own expression in AVK and DVA neurons. Top panel shows *hlh-15* locus. The whole upstream intergenic region (from ATG to −777bp) was fused to *gfp* to generate *wwEx42* (Grove *et al*. 2009). Representative pictures and quantification showing expression of *hlh-15/NHLH* promoter fusion (*wwEx42*) in *hlh-15/NHLH* mutants. Animals were scored at the L1 (middle panel) and young adult stage (bottom panel). Statistical analysis was performed using Fisher’s exact test. N is indicated within each bar and represents number of animals scored.

AVK and DVA are also among the very few neuron classes that continuously express the E/Da homolog HLH-2 throughout postembryonic life (**Fig.1A**)(Krause *et al*. 1997; Masoudi *et al*. 2022). This indicates that HLH-15/NHLH may, like its vertebrate homolog (Uittenbogaard *et al*. 1999), heterodimerize with the E protein HLH-2, a notion supported by HLH-15::HLH-2 protein interactions in yeast 2-hybrid assays (Grove *et al*. 2009).

Continuous expression of transcription factors throughout the life of a cell are often reflection of transcriptional autoregulation (Hobert 2011; Leyva-DIAZ AND HOBERT 2019). To ask whether *hlh-15/NHLH* autoregulates, we utilized a previously described transcriptional reporter of the *hlh-15/NHLH* locus in which its intergenic 5’ promoter region was fused to *gfp* (Grove et al. 2009). This reporter recapitulates continuous expression in the AVK and DVA neurons (**Fig.7D**). In an *hlh-15/NHLH* null mutant background, expression of the reporter is initially normally observed in AVK and DVA, but it fades during postembryonic life to result in complete absence at the adult stage (**Fig.7D**). This observation demonstrates autoregulation of the *hlh-15/NHLH* locus.

### The peptidergic AVK neuron class fails to differentiate properly in *hlh-15/NHLH* mutant animals

We used two mutant alleles to analyze *hlh-15/NHLH* function: (1) a deletion mutant, *tm1824,* generated by the NBRP knockout consortium, which deletes most of the locus, including most of its bHLH domain and (2) a complete locus deletion, *ot1389*, that we generated by CRISPR/Cas9 genome engineering (**Fig.1C**). Animals carrying either allele are fully viable and display no obvious morphological abnormalities. Unlike their vertebrate counterparts (Good and Braun 2013), *hlh-15/NHLH* mutants display no obvious fertility defects.

We assessed the potential function of *hlh-15/NHLH* in controlling fate specification of the AVK and DVA neurons by first using NeuroPAL, the marker transgene in which all neurons express specific codes of cell fate markers (Yemini *et al*. 2021). In NeuroPAL, the AVK neurons are marked with the *flp-1* neuropeptide gene and DVA is marked with the *nlp-12* neuropeptide and the two ionotropic glutamate receptors *nmr-1* and *glr-1.* NeuroPAL colors in *hlh-15/NHLH* mutants indicate a loss of the AVK color code (*flp-1* expression), but no effect on the color code of the DVA neuron (**Fig.8A**). We confirmed the proper differentiation of DVA in *hlh-15/NHLH* mutants using another DVA fate marker, a reporter allele for the acetylcholine vesicular transporter, *unc-17/VAChT* whose expression we found to be unaffected (**Fig.8B**).

**Fig. 8:**
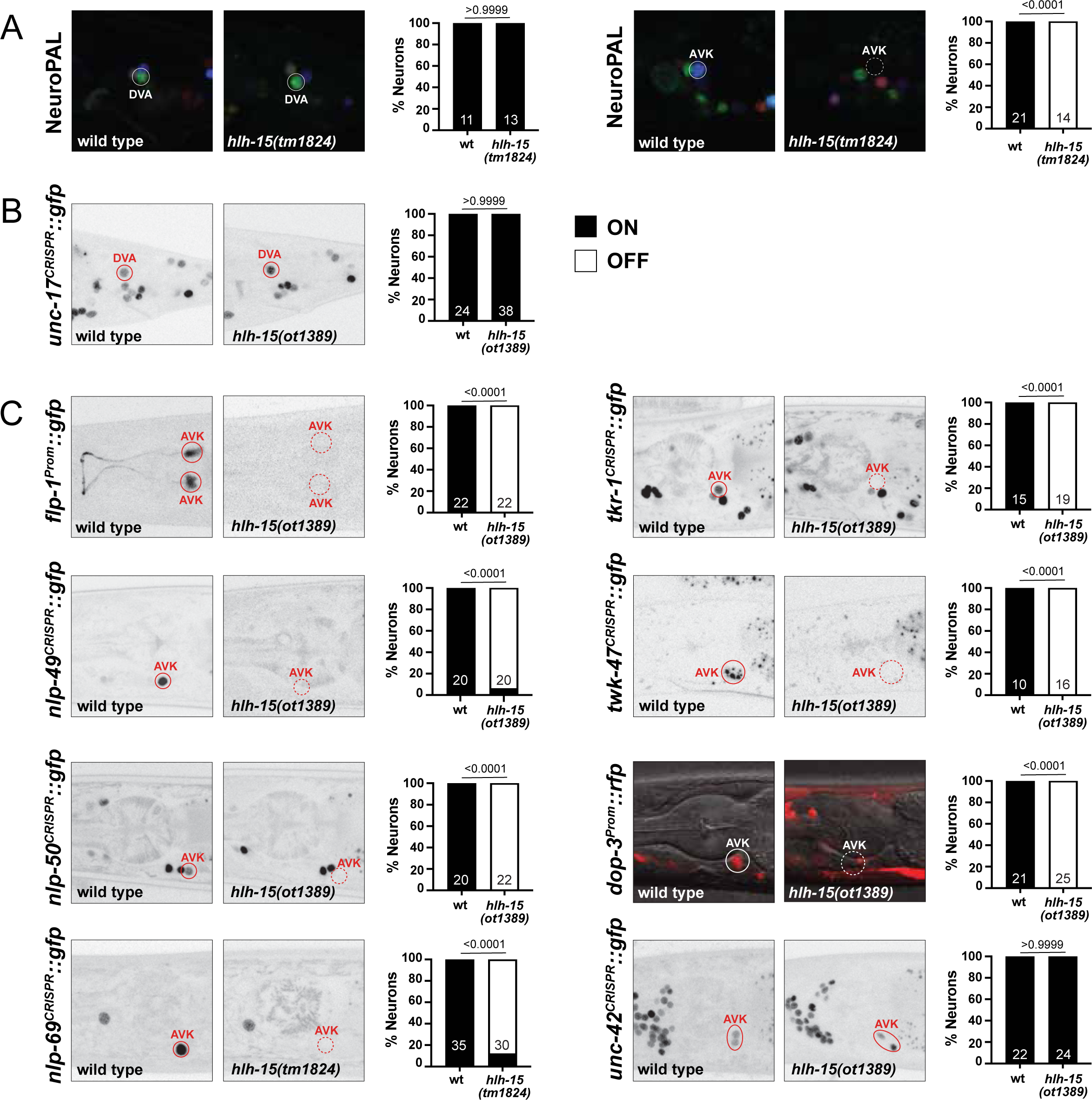
*hlh-15/NHLH* gene affects the differentiation of the AVK interneuron. **A:** Representative pictures and quantification showing unaffected NeuroPAL (*otIs669*) coloring in DVA (left panel) and loss of NeuroPAL (*otIs669*) coloring in AVK (right panel) in *hlh-15/NHLH* mutants. Animals were scored at the young adult stage. Statistical analysis was performed using Fisher’s exact test. N is indicated within each bar and represents number of animals scored. **B:** Representative pictures and quantification showing *unc-17* expression in DVA is unaffected in *hlh-15/NHLH* mutants. Reporter gene used is CRISPR/Cas9-engineered reporter allele *unc-17* (*syb4491*). Animals were scored at the young adult stage. Statistical analysis was performed using Fisher’s exact test. N is indicated within each bar and represents number of animals scored. **C:** Representative pictures and quantification showing loss of cell fate markers in *hlh-15/NHLH* mutants. Reporter genes used are transgenes *flp-1*(*bwIs2*), dop-3(*vsIs33*) and CRISPR/Cas9-engineered reporter alleles *nlp-49*(*syb3320*), *nlp-50*(*syb6148*), *nlp-69*(*syb4512*), *tkr-1*(*syb4595*), *twk-47*(*bab385*) and *unc-42*(*ot986*). Animals were scored at the young adult stage. Statistical analysis was performed using Fisher’s exact test. N is indicated within each bar and represents number of animals scored.

We confirmed the AVK differentiation defects using another transgene to measure *flp-1* expression, *bwIs2* (Much et al. 2000)(**Fig.8C**). Given that AVK is a unique neuropeptidergic hub interneuron, receiving and sending a multitude of neuropeptidergic signals (Ripoli-SANCHEZ *et al*. 2022), we further probed the acquisition of AVK’s neuropeptidergic identity in *hlh-15/NHLH* mutant animals.

Specifically, we probed the peptidergic identity of AVK by generating reporter alleles for three additional neuropeptides, predicted by scRNAseq data to be expressed in AVK, *nlp-49, nlp-50* and *nlp-69* (Taylor et al. 2021). *nlp-49* was previously also reported to be required in AVK to control motor behavior (Chew *et al*. 2018; Oranth *et al*. 2018) and is, next to *flp-1*, the neuropeptide that is most selectively expressed in AVK (Taylor et al. 2021). We CRISPR/Cas9-engineered reporter alleles for all three genes and found that expression of *nlp-49* in AVK is completely lost in *hlh-15/NHLH* mutant animals, while expression in other *nlp-49(+)* neurons is unaffected (**Fig.8C**). Similarly, expression of the *nlp-50* and *nlp-69* reporter alleles is also selectively lost in the AVK neurons of *hlh-15/NHLH* mutants (**Fig.8C**).

Apart from transmitting neuropeptidergic signals, AVK also expresses a multitude of neuropeptidergic receptors (Taylor *et al*. 2021; Ripoli-SANCHEZ *et al*. 2022). We examined one of them, the tachykinin receptor *tkr-1,* expressed in AVK and many other head neurons (Ripoli-SANCHEZ *et al*. 2022), and found its expression in AVK to be disrupted in *hlh-15/NHLH* mutant animals (**Fig.8C**). AVK also receives dopaminergic signals via the AVK-expressed dopamine receptor *dop-3* (Ji *et al*. 2023), and we find *dop-3* expression to also be disrupted in *hlh-15/NHLH* mutant animals (**Fig.8C**).

The nervous system-wide scRNA atlas of *C. elegans* predicts another unique identity marker of the AVK neurons, a two pore TWIK-type potassium channel, *twk-47* (Taylor et al. 2021). Its selective expression in AVK was recently validated with a promoter-based transgene construct (Lorenzo *et al*. 2020) and is further validated with a CRISPR/Cas9-genome engineered, wrmScarlet-based reporter allele (kindly provided by T. Boulin). We find that expression of this *twk-47* reporter allele is also completely eliminated from the AVK neurons of *hlh-15/NHLH* mutants (**Fig.8C**).

Contrasting effects on neuron type-specific terminal effector genes, we found that *hlh-15/NHLH* null mutant animals show normal expression of pan-neuronal genes, as well as normal expression of a previously described terminal selector of AVK differentiation, the *unc-42* homeobox gene (**Fig.8C**). These observations indicate that (a) *hlh-15/NHLH* is not acting as a proneural gene to determine the generation of the AVK neurons and (b) that *hlh-15/NHLH* may work together, perhaps in a heteromeric, combinatorial manner, with *unc-42* as a terminal selector of AVK identity (see Discussion).

### Proprioceptive defects in *hlh-15/NHLH* mutant animals

The AVK neurons were previously shown to act downstream of dopaminergic neurons to control proprioceptive behavior, a process mediated by the AVK-released and *hlh-15*-controlled *flp-1* gene (Ji *et al*. 2023). Given that *hlh-15/NHLH* controls *flp-1* expression in AVK (the only *C. elegans* neuron that expresses *flp-1*), we predicted that *hlh-15/NHLH* mutants might display similar defects in proprioceptive behavior. We tested this prediction by measuring the AVK- and *flp-1-*dependent compensatory curvature response in animals carrying either of the two available *hlh-15/NHLH* deletion alleles and found a severe reduction of the response, mimicking the genetic ablation of AVK (**Fig.9A**).

**Fig. 9:**
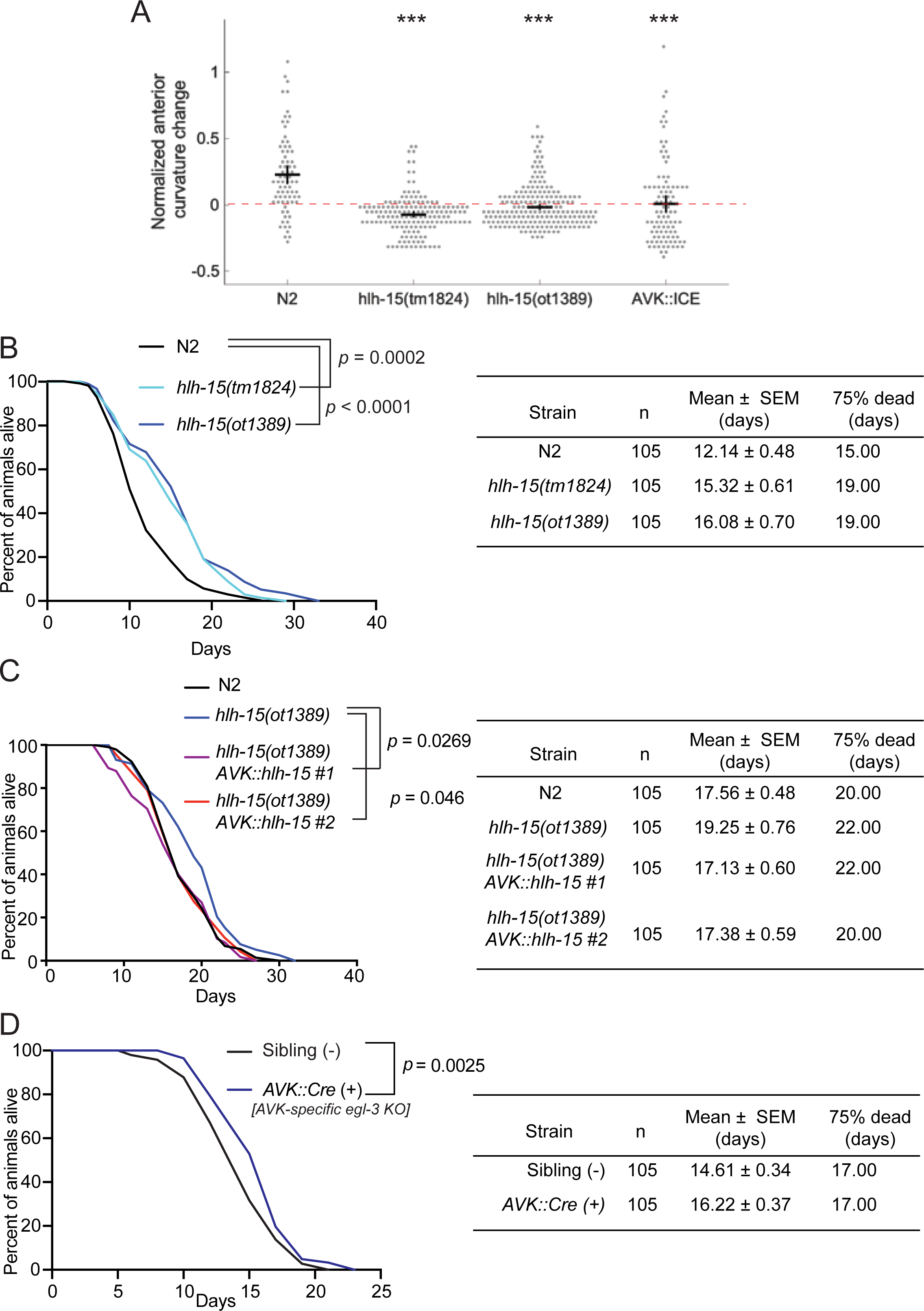
Behavioral and physiological analysis of *hlh-15/NHLH* mutants. **A:** *hlh-15(tm1824)* and *hlh-15(ot1389)* null mutants display defects in their compensatory curvature response (CCR)(Ji *et al*. 2023). These defects are similar to those observed upon genetic ablation of AVK using a transgenic array, *zxIs28[pflp-1(trc)::ICE; pmyo-2::mCherry],* in which the ICE caspase was driven by an AVK-specific *flp-1* promoter (Oranth *et al*. 2018). N ≥ 10 animals per condition. Error bars indicate mean ± SEM. ***P < 0.001, Dunnett’s multiple comparison tests. **B:** *hlh-15(tm1824)* and *hlh-15(ot1389)* null mutants show increased lifespan. Survival plot (left) and mean and 75% mortality lifespan (right) of N2, *hlh-15(tm1824)*, and *hlh-15(ot1389)* animals. A repeat of these lifespan assays is shown in Fig.S4. **C:** Lifespan extension defects of *hlh-15(ot1389)* mutants are rescued by expressing *hlh-15/NHLH* selectively in the AVK neurons (*flp-1^prom^::hlh-15).* Survival plot (left) and mean and 75% mortality lifespan (right) of N2, *hlh-15(ot1389)* and two independent transgenic lines (*otEx8247* and *otEx8248)* of *hlh-15(ot1389)* animals expressing genomic *hlh-15* DNA specifically in AVK. A repeat of these lifespan assays are shown in Fig.S4. This assay could not be performed with the AVK::ICE lines because animals had a sick appearance. **D:** Removal of neuropeptide processing in AVK neurons extends lifespan. Survival plot (left) and mean and 75% mortality lifespan (right) of *egl-3(nu1711)* animals with the transgene (*kyEx6532*) or without the transgene (siblings) that expresses Cre recombinase specifically in AVK. For panels B, C and D, statistics were performed using Log-Rank Test followed by Bonferroni corrections. N = 105 for each strain.

### *hlh-15/NHLH* acts in the AVK interneuron class to regulate life span

RNAi knockdown of *hlh-15,* as well as knock-down of its presumptive, class I heterodimerization partner *hlh-2,* has previously been reported to extend life span of *C. elegans* (Mansfeld *et al*. 2015; Rozanov *et al*. 2020). We found that the reported lifespan extension effects of *hlh-15(RNAi)* animals are recapitulated in animals that carry either of the two independent deletion alleles of the *hlh-15/NHLH* locus (**Fig.9B**).

The previous RNAi-based analysis of *hlh-15/NHLH* (Mansfeld *et al*. 2015) had not considered the potential cellular focus of action of *hlh-15/NHLH* for lifespan control. Since loss of *hlh-15/NHLH* only affects the differentiation of the AVK neuron, and not the only other neuron (DVA) in which *hlh-15/NHLH* is expressed in, we tested the possibility that *hlh-15/NHLH* acts in AVK to control animal lifespan. Specifically, we asked whether the extended lifespan of *hlh-15/NHLH* is restored to normal wildtype-like lifespan by resupplying *hlh-15/NHLH* selectively in the AVK interneurons. We found that a transgenic animals that re-express *hlh-15/NHLH* from an AVK-specific promoter in an *hlh-15/NHLH* null mutant background indeed displayed a rescue of the lifespan extension effect of *hlh-15* mutants (**Fig.9C; Fig.S4**), indicating that a single interneuron has the capacity to control animal life span.

Previous work has shown lifespan extension of animals in which the expression of an enzyme involved in branched-chain amino acid metabolism, *bcat-1 (*<u>b</u>ranched-<u>c</u>hain <u>a</u>mino acid transferase-1) was reduced by RNAi (Mansfeld *et al*. 2015). Moreover, animal-wide RT-qPCR analysis suggested that levels of *bcat-1* transcripts are reduced in *hlh-15(RNAi)* animals (Mansfeld *et al*. 2015). A putative HLH-15/NHLH binding site in the *bcat-1* promoter made these authors suggest that HLH-15/NHLH directly controls *bcat-1* expression. We sought to independently probe the potential link of *hlh-15/NHLH* and *bcat-1* with alternative approaches. To this end, we first tagged the *bcat-1* locus with *gfp*, using CRISPR/Cas9 genome engineering. As expected from a metabolic enzyme, we found very broad (albeit not entirely ubiquitous) and robust expression of *bcat-1* across all major tissue types (**Fig.S5**). However, in a *hlh-15/NHLH* null mutant background, *bcat-1* expression appeared unchanged, in both young or aged adult animals (**Fig.S5**). While this experiment does not entirely rule out that the lifespan extending effects of *hlh-15/NHLH* may be mediated by *bcat-1*, it provides a first indication that *hlh-15/NHLH* and AVK may act through other means to control animal lifespan.

One set of possible life-span modulating effector genes of *hlh-15/NHLH* are neuropeptides secreted by AVK. Based on scRNAseq data, as well as reporter gene data, AVK shows highly selective expression of at least eight neuropeptide-encoding genes, some of which deeply conserved throughout animal phylogeny (Chew *et al*. 2018; Taylor *et al*. 2021; Beets *et al*. 2022; Ripoll-SANCHEZ *et al*. 2022). As demonstrated above, the expression of at least some, if not all AVK-expressed neuropeptides is under *hlh-15/NHLH* control (**Fig.8C**). Rather than venturing into testing life-span effects of individual neuropeptide-encoding gene genes, we eliminated all neuropeptide processing selectively in the AVK neuron, using a conditional, floxed allele of the *egl-3* proprotein convertase required for neuropeptide processing (Kass *et al*. 2001; Marquina-SOLIS *et al*. 2024). We found that animals in which we removed *egl-3* in AVK also showed an increased lifespan (**Fig.9D**). We conclude that the lifespan-extending effect of *hlh-15/NHLH* can be explained via *hlh-15/NHLH* controlling the neuropeptidergic phenotype of AVK and, more generally, that a single interneuron has the capacity to influence the lifespan of an animal.

## DISCUSSION

### Revising expression patterns of conserved bHLH proteins in *C. elegans*

We have discovered functions of several, phylogenetically conserved bHLH proteins during terminal differentiation of postmitotic neurons, thereby expanding our understanding of bHLH gene function in the nervous system. The three clustered bHLH genes *hlh-17/31/32* were originally considered orthologs of Olig proteins, which are important regulators of glia and motor neuron differentiation in vertebrates (Lu *et al*. 2002; Takebayashi *et al*. 2002; Zhou and Anderson 2002). However, by sequence homology, *hlh-17/31/32* are the orthologs of the vertebrate bHLHe4 and bHLHe5 proteins rather than the closely related vertebrate Olig proteins. Unlike the vertebrate Olig proteins, vertebrate bHLHe4/e5 have no reported function in glia cells, but rather act in neuronal differentiation in select neuron classes in the retina, telencephalon and spinal cord (Bramblett *et al*. 2004; Feng *et al*. 2006; Joshi *et al*. 2008; Ross *et al*. 2010; Skaggs *et al*. 2011; Oyallon *et al*. 2012; Ross *et al*. 2012; Huang *et al*. 2014).

The *Drosophila* bHLHe4/e5 homolog also functions in motor neurons rather than glia (Oyallon *et al*. 2012). Similarly, we find here that the *C. elegans* bHLHe4/5 orthologs HLH-17 and HLH-32 are expressed and function in neurons, including motor neurons, but not glia. We cannot exclude that HLH-32/17/31 proteins are expressed below the levels of reporter allele-based detectability in the CEPsh glia cells. However, given the robust transcription of the *hlh-17/31/32* genes in CEPsh glia (based on scRNAseq and promoter-fusion reporters), we consider posttranscriptional repression the most parsimonious explanation. Such posttranscriptional regulation is evident upon comparison of mRNA transcripts and homeodomain protein expression in several distinct neuron types (Reilly *et al*. 2020; Taylor *et al*. 2021).

We also found that the expression pattern of a reporter allele of the PTF1a homolog *hlh-13* is different from the previously reported expression of a multicopy *hlh-13/PTF1a* reporter transgene (Liachko *et al*. 2009; Bou DIB *et al*. 2014). However, in this case, we do not infer posttranscriptional regulation as a source for differences, since our reporter allele data is consistent with scRNAseq data. We rather consider previous reporter genes to either produce incorrect expression and/or the sites of expression have been incorrectly identified, a problem remedied by our usage of the neuronal landmark strain NeuroPAL.

Lastly, NeuroPAL has also been instrumental in defining the proper identity of *hlh-15/NHLH* expressing cells, which had only been incompletely achieved with previous reporter transgenes (Grove *et al*. 2009). Taken together, our expression pattern analysis of these 5 conserved bHLH genes illustrated the importance of using the best possible tools (reporter alleles and cell ID tools) to properly infer protein expression patterns.

### Function of bHLH genes in neuronal identity specification

We discovered terminal neuronal differentiation defects for each of the bHLH subfamilies that we examined in this study. The bHLHe4/5 orthologs *hlh-17* and *hlh-32* function as “terminal subtype selectors” that diversify motor neuron subclass identity in the retrovesicular ganglion that harbors neurons akin to vertebrate branchial motor neurons. In another neuron class, AUA, which is also potentially involved in motor control (Yemini *et al*. 2021), *hlh-17* and *hlh-32* affect select, but not all aspects of the differentiation program. *hlh-17* and *hlh-32* function in motor neuron differentiation is reminiscent of the function of *Drosophila* and vertebrate homologs of these genes, which play a role in various aspects of motor neuron specification (Lu *et al*. 2002; Zannino and Appel 2009; Oyallon *et al*. 2012; Alfano and Studer 2013), but we added here some intriguing granular detail to such motor neuron function. While anatomical reconstructions (White *et al*. 1986) and, more recently, scRNAseq analysis (Taylor *et al*. 2021; Smith *et al*. 2024), has revealed that the most anteriorly located B-type motor neurons within the RVG are morphologically and molecular distinct from other B-type motor neurons along the ventral nerve cord, no molecular mechanisms for their subclass diversification have been previously identified. We found that *hlh-17* and *hlh-32* do not affect the expression of features shared by B-type neuron types, but instead promote the expression of features that are characteristic of one subtype (VB2), while repressing features characteristic of another subtype (VB1). Generally, all motor neuron markers, including B-type motor neurons markers and at least some, if not all subtype-specific markers described here, are directly controlled by the terminal selector *unc-3,* an EBF-type transcription factor (Kratsios *et al*. 2011; Kerk *et al*. 2017). Akin to other subtype diversifiers (e.g. BNC-1)(Kerk *et al*. 2017), we propose that *hlh-17/32* act as subtype diversifiers to either directly or indirectly promote or antagonize the ability of UNC-3 to turn on select target genes.

*hlh-13/PTF1a* and *hlh-15/NHLH* appear to act in a canonical terminal selector-type manner in two different neuron classes, *hlh-13/PTF1a* in the octopaminergic RIC interneurons class and *hlh-15/NHLH* in the peptidergic AVK interneuron class. In both neuron classes, loss of the respective bHLH gene affects the expression of all markers tested, but they do not affect the generation of the neurons and they are continuously expressed throughout the life of the neuron to possible maintain their identity. Such continuous expression is likely ensured by transcriptional autoregulation, as we have explicitly shown for *hlh-15/NHLH*.

The terminal selector role of *hlh-13/PTF1a* and *hlh-15/NHLH* appears similar to the role of the ASC-type *hlh-4* bHLH as a validated terminal selector of the ADL sensory neuron class, where HLH-4 operates, likely in conjunction with the common heterodimerization partner, the E/Da-homolog HLH-2, to co-regulate scores of terminal effector genes through direct binding to E-box motifs (Masoudi *et al*. 2018). In both the AVK and RIC neuron classes, the HLH-15/NHLH and HLH-13/PTF1A protein, respectively, are also likely to heterodimerize with the common class I E/Da HLH-2 protein, based on: (a) co-expression of HLH-13/PTF1A and HLH-15/NHLH with HLH-2, which is expressed in 8 neuron classes, including the AVK and DVA neurons (ADL, ASH, PHA, PHB, RIC, AVK, DVA, PVN)(Krause *et al*. 1997; Masoudi *et al*. 2022); (b) physical interaction of HLH-15::HLH-2 and HLH-13::HLH-2 tested in protein-protein interaction assays (Grove *et al*. 2009); (c) similar interactions of the vertebrate homolog of HLH-13 (PTF1a) and HLH-15 (NHLH1/2) with vertebrate E-proteins (Uittenbogaard *et al*. 1999; Rose *et al*. 2001). *hlh-2* is not expressed in the *hlh-17/31/32-*expressing AUA/DB2/VB2 neurons, but this is consistent with Olig-related proteins acting as strong homodimers (Li and Richardson 2008). It is notable that for some of the neurons that express the common E/Daughterless bHLH component HLH-2 postembryonically, no expression of other class A bHLH proteins is observed (**Fig.1A**). This indicates that in these neurons (e.g. ASH or PHB), *hlh-2* may act as a homodimer, as it does in other non-neuronal cells (Sallee and Greenwald 2015). A role of a terminal selector HLH-2 homodimer is conceivable because E-box motifs are enriched in the ASH and PHB terminal gene batteries (Taylor *et al*. 2021).

Vertebrate homologs of *hlh-13* and *hlh-15* may have maintained their function as regulators of neuronal differentiation. The vertebrate bHLH PTF1a and its paralogue NATO3 have multiple functions both early and late during neuronal development (Ono *et al*. 2010; Jin and Xiang 2019). The vertebrate *hlh-15* homologs NHLH1 and NHLH2 control the proper differentiation, and perhaps also maintenance, of several peptidergic hormone-producing cells in the hypothalamus (Kruger *et al*. 2004; Good and Braun 2013; Leon *et al*. 2021), reminiscent of the role of *hlh-15/NHLH* in controlling the identity of the peptidergic AVK neuron. Roles in initiating and maintaining terminal differentiation programs as putative terminal selectors have, however, not yet been explicitly investigated for those vertebrate homologs, something that should be tested through temporally controlled removal of vertebrate homologs specifically during or after terminal differentiation.

### Discovery of a lifespan-controlling peptidergic hub neuron

The very restricted expression of *hlh-15/NHLH* in exclusively two neuron classes, combined with its striking function in controlling animal lifespan led us to discover a function of a single interneuron class, AVK, in controlling lifespan of the animal. The impact of the *C. elegans* nervous system on animal lifespan has long been appreciated (Apfeld and Kenyon 1999; Alcedo and Kenyon 2004), but has largely been restricted to the sensory periphery, which releases distinct types of neuropeptides, mostly insulins, but also other types of neuropeptides to modulate animal ageing (Artan *et al*. 2016; Chen *et al*. 2016; Zhang *et al*. 2018). We discover here a different type of neuron that controls lifespan, the AVK interneurons.

The AVK neuron class has been shown to regulate responses to various physiological states to control locomotion and behavior (Hums *et al*. 2016; Chew *et al*. 2018; Oranth *et al*. 2018; Ji *et al*. 2023; Marquina-SOLIS *et al*. 2024), but AVK has not previously been implicated in lifespan control. Moreover, AVK displays fundamental differences to neurons previously implicated in lifespan control. It is neither a sensory neuron, nor a main postsynaptic target of sensory neurons (White *et al*. 1986). Its expression of a large number of neuropeptide receptors as well as of neuropeptides that act through receptors distributed throughout the animal nervous system has revealed AVK to be a central neuropeptidergic hub neuron (Ripoll-SANCHEZ *et al*. 2023) that likely controls internal states of the animal. We hypothesize that AVK may serve as an integrator of various internal states, resulting in the release of neuropeptides that control animal lifespan.

Both in terms of its peptidergic nature and also its developmental specification, AVK is conceptually remarkably similar to peptidergic neurons in the mammalian hypothalamus. Peptidergic hypothalamic neurons control internal states of mammals and have been implicated in lifespan control as well (Flurkey *et al*. 2001; Zhang *et al*. 2013; Kim and Choe 2019). Strikingly, the mammalian homologs of the AVK-specifying *hlh-15* gene, NHLH1 and NHLH2, control the proper specification of several classes of peptidergic neurons of the hypothalamus (Kruger *et al*. 2004; Good and Braun 2013; Leon *et al*. 2021). It will be intriguing to define with better resolution the exact type of hypothalamic neurons that continuously express and require NHLH1/2 during adulthood as this may help to more precisely determine – in analogy to HLH-15/NHLH expression and function – the nature of the neurons that may control lifespan in mammals.

Intriguingly, the presently completely uncharacterized sole homolog of NHLH1/2 and HLH-15 in *Drosophila*, the HLH4C gene, appears to be expressed in the pars intercerebralis, which is considered to be the hypothalamus in *Drosophila* (De VELASCO et al. 2007).

### bHLH genes and homeobox gene as regulators of neuronal identity

The bHLH proteins that we have characterized here are unlikely to act in isolation but can rather be expected to act in combination with other transcription factors to control terminal neuron differentiation as terminal selectors. Their most likely partners are homeodomain transcription factors. Each neuron class in *C. elegans* is defined by a unique combination of homeodomain proteins and mutant analysis has corroborated their widespread deployment as drivers of terminal neuron differentiation programs. Based on previous mutant analysis, HLH-15/NHLH (likely in combination with HLH-2) may cooperate with the Prop1-type homeodomain protein UNC-42 in the AVK neurons, where UNC-42 had previously shown to display similar differentiation defects to those that we have described here (Much *et al*. 2000; Berghoff *et al*. 2021). Similarly, in the octopaminergic RIC neurons, HLH-13/PTF1A likely acts together with the Pbx-type homeodomain protein *unc-62,* whose loss also results in RIC differentiation defects (Reilly *et al*. 2022). And even though *hlh-17/31/32* may only control aspects of AUA differentiation, a potential cofactor is yet another homeobox gene, the POU homeobox gene *ceh-6*, which controls AUA differentiation (Serrano-SAIZ *et al*. 2013). Direct partnerships between bHLH and homeodomain proteins have been observed in other systems as well (Poulin *et al*. 2000; Sun *et al*. 2001; Wang and Harris 2005).

Taking a step back and comparing the relative functional deployment of the 42 bHLH family members and 102 homeobox genes in inducing and maintaining terminal differentiation programs in *C. elegans*, it is apparent that homeobox genes cover the differentiating nervous system much more broadly than bHLH factors do. Mutant analyses of many bHLH and homeobox genes support the notion that with notable exceptions (such as those described here), bHLH genes tend to be more involved in early patterning events, while homeobox genes appear to be biased toward controlling terminal differentiation events in the nervous system. It will be fascinating to see whether this is an evolutionary conserved theme.

## ACKNOWLEDGEMENTS

We thank Chi Chen for *C. elegans* microinjection, Shohei Mitani at Tokyo Women’s Medical University School of Medicine, Tokyo for tm alleles, Surojit Sural for advice on lifespan assays and members of the Hobert lab for comments on the manuscript. Some strains were provided by the CGC, which is funded by NIH Office of Research Infrastructure Programs (P40 OD010440). We thank Thomas Boulin for providing the *twk-47* reporter allele which was generated by SEGiCel (SFR Santé Lyon Est CNRS UAR 3453, Lyon, France). This work was supported by NIH R01NS110391 and NIH R01NS039996.

## AUTHOR CONTRIBUTIONS

GRA performed all experiments related to *hlh-17, hlh-31* and *hlh-32*.

BV and JE performed cell fate analysis of *hlh-13* and *hlh-15* mutants as well as expression pattern analysis of *hlh-13* and *hlh-15.* GV analyzed a subset of markers. HJ performed lifespan assays and fat staining experiments under supervision of CPL.

CPL generated reagents and transgenic lines.

HJ performed locomotor and CCR analyses under supervision of CFY.

OH initiated and supervised the project and wrote the manuscript, which was read and co-edited by all co-authors.

## SUPPLEMENT FIGURES

**Fig. S1:**
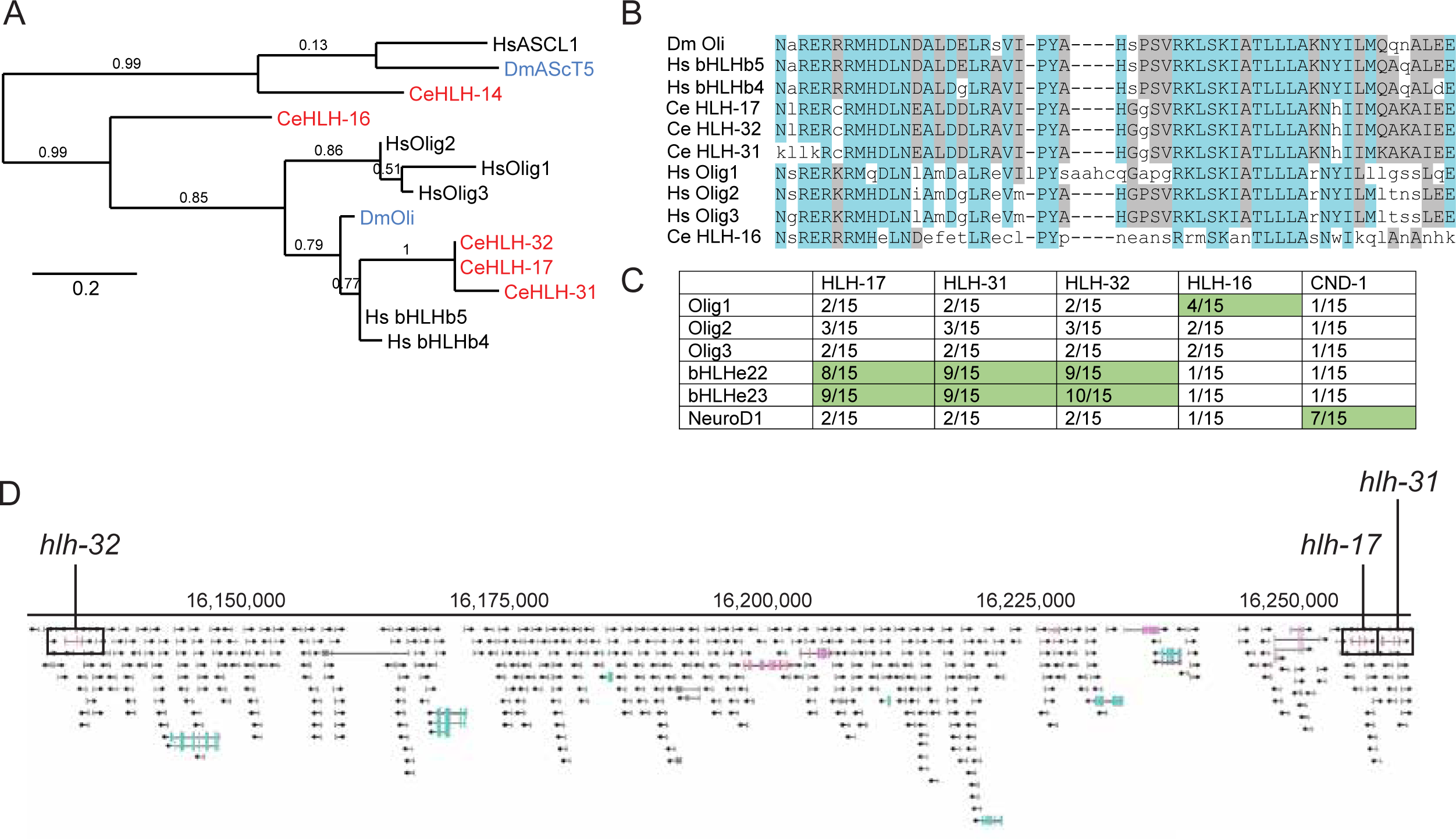
Molecular phylogeny of the Olig/bHLHb4/b5 subfamily. **A:** Phylogenetic relationship of *C. elegans* and human Olig/bHLHb4/b5 members. Tree generated at phylogeny.fr (Dereeper *et al*. 2008) with default parameters. **B:** Protein sequence alignment of Olig/bHLHb4/b5 members across phylogeny. Created at phylogeny.fr by MUSCLE. *Similar residues are colored as the most conserved one (according to BLOSUM62). Colores indicate average BLOSUM62 score: blue 1.5, low 0.5.* **C:** Tabular DiOPT scores of Olig/bHLHb4/b5 members as provided by Marrvel (Wang *et al*. 2017). One NeuroD homolog is provided as an “outgroup”. **D:** *hlh-32, hlh-17, and hlh-31* are located close to each other in a region of C. elegans chromosome IV that is replete with small RNAs. From the genome browser of WormBase.

**Fig. S2:**
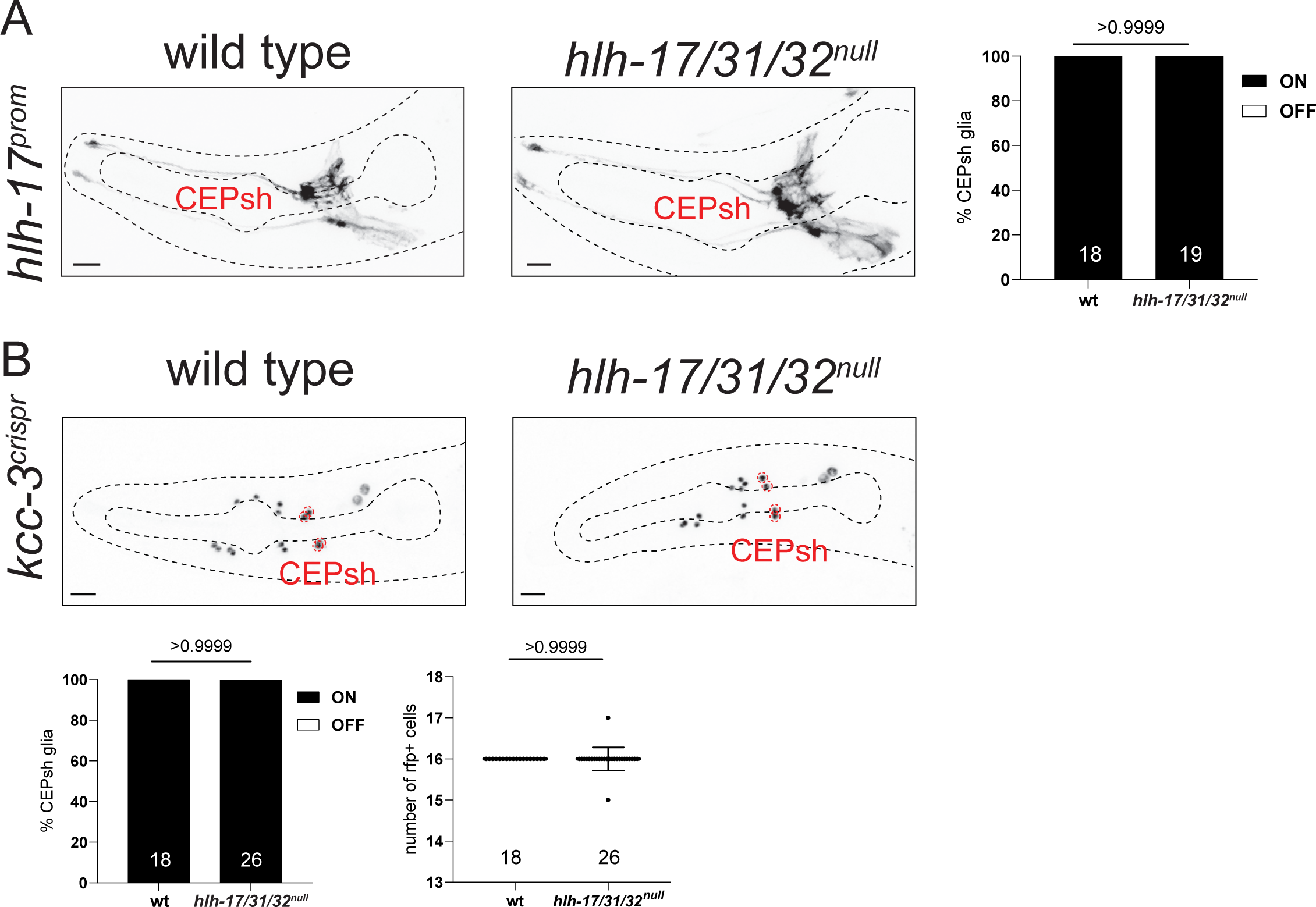
*hlh-17/31/32^null^* animals show not measurable effects on CEPsh morphology or marker expression. **A:** *hlh-17/31/32^null^* animals show no defects in in the expression of the CEPsh marker *irIs67 (hlh-17prom::gfp).* CEPsh morphology is also unaffected. Representative images of wild type and mutant animals are shown with 10 μm scale bar. Number of animals scored are within each bar. P-values were calculated using Fisher’s exact test. **B:** *hlh-17/31/32^null^* animals show no defects in the expression of glial marker *kcc-3(syb4430).* Expression in CEPsh is still clearly identifiable. Expression in other glia was also unaffected as indicated by counting the number of *kcc-3-*expressing cells. Representative images of wild type and mutant animals are shown with 10 μm scale bar. Number of animals scored are shown within each bar. Statistical analysis for CEPsh expression was done using Fisher’s exact test, while that for counting *kcc-3*-expressing cells was performed using unpaired t-test. Error bars for the scatter plot indicate standard deviation of the mean.

**Fig. S3:**
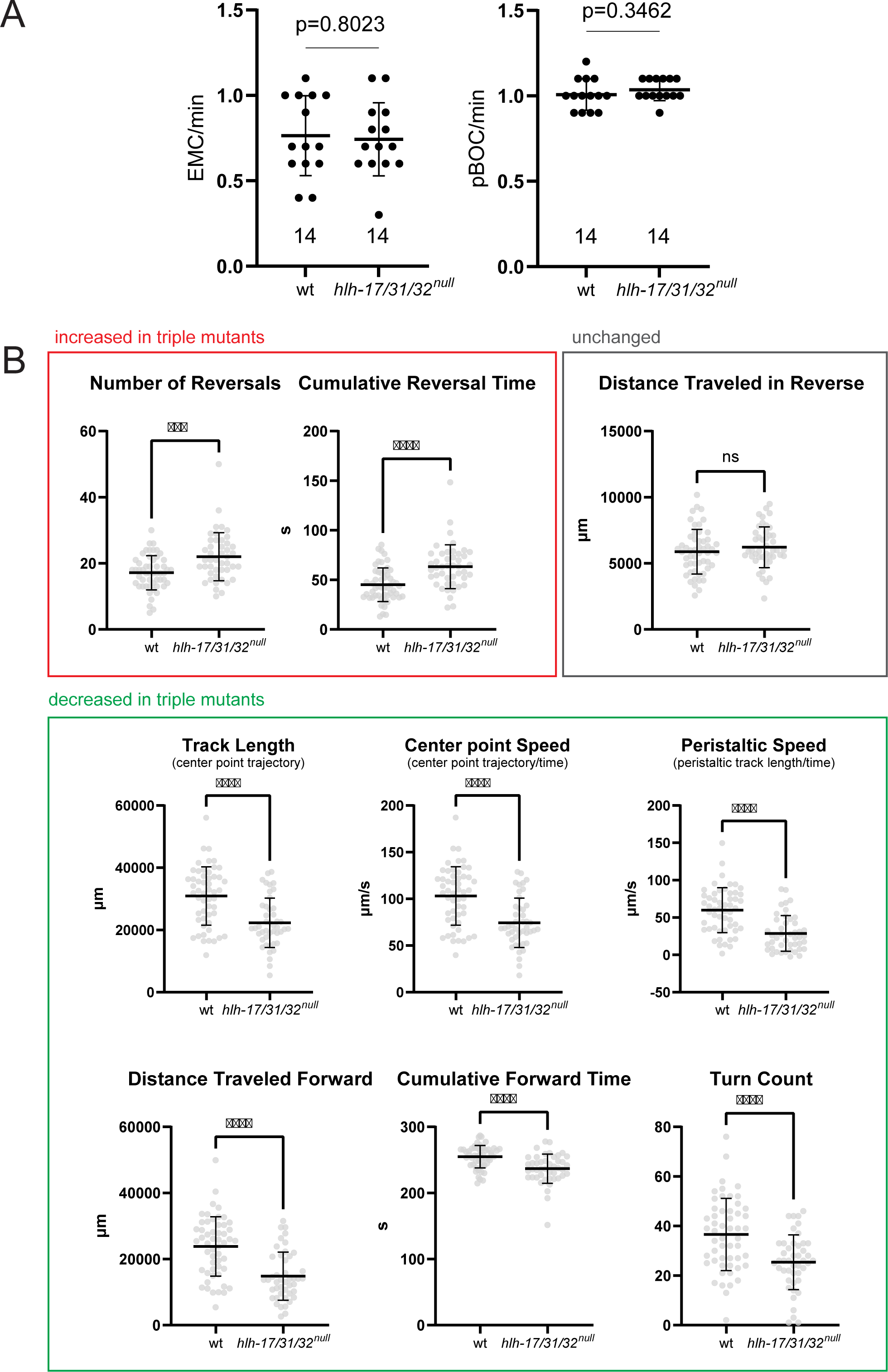
Behavioral analyses of *hlh-17/31/32^null^ animals*. **A:** *hlh-17/31/32^null^* animals do not display defects in the defecation motor program, contrary to a previous report using a different *hlh-17* single mutant allele (Bowles and Johnson 2021). Individual points represent measurements of individual worms. Statistical analysis was performed using unpaired t-test. Error bars indicate standard deviation of the mean. ns: not significant. **B:** Worm tracking of *hlh-17/31/32^null^* animals reveal locomotory defects. Individual points represent measurements of individual worms. Data are pooled from three independent experiments. Statistical analysis was performed using unpaired t-test. Error bars indicate standard deviation of the mean. ****p ≤ 0.0001, ***p ≤ 0.001, ns: not significant. N=53 for wild type, N=45 for *hlh-17/31/32^null^*.

**Fig. S4:**
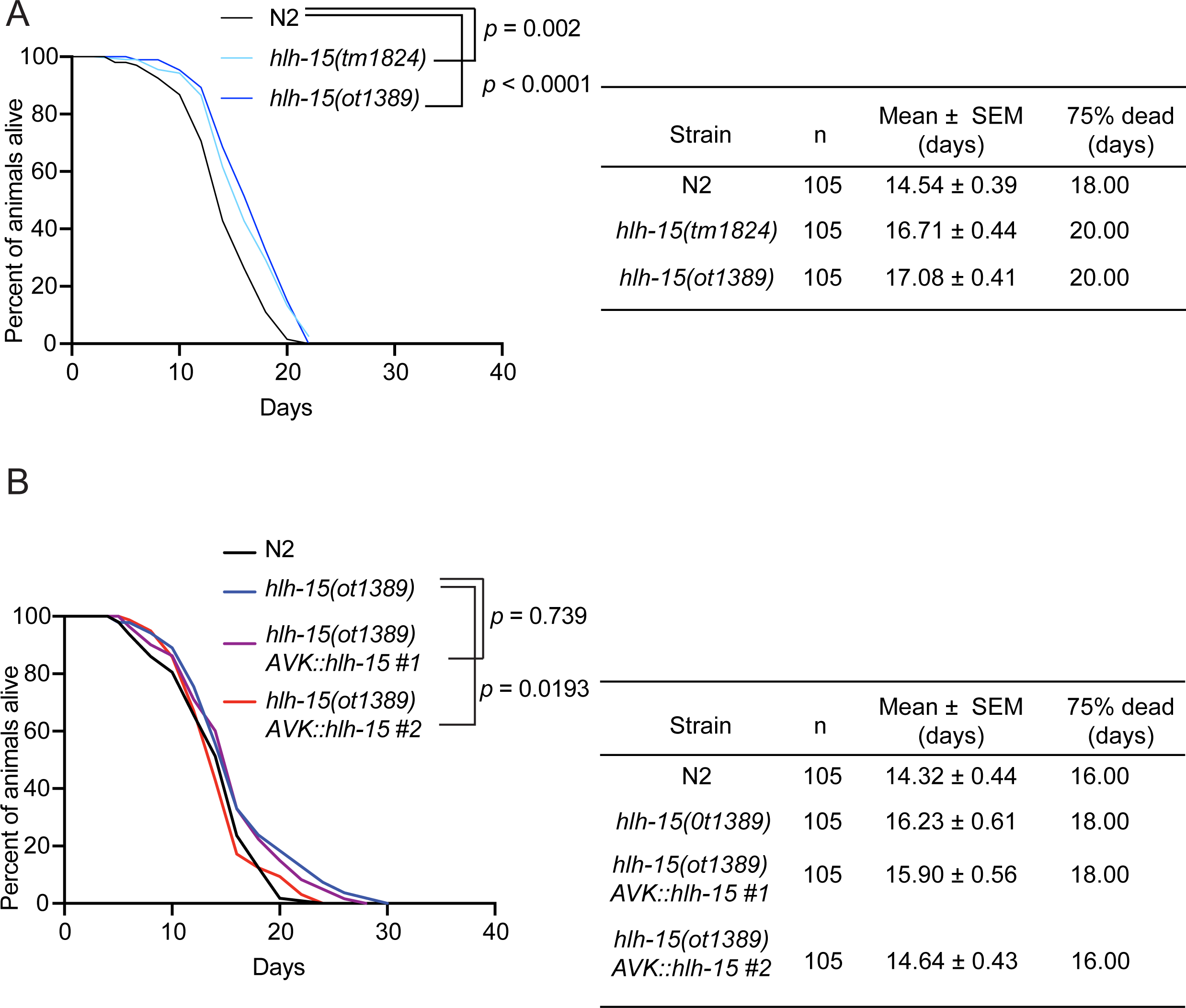
Replication of *hlh-15/NHLH* lifespan assays. **A:** Replication of the experiment in Fig.9B. **B:** Replication of the experiment in Fig.9C.

**Fig. S5:**
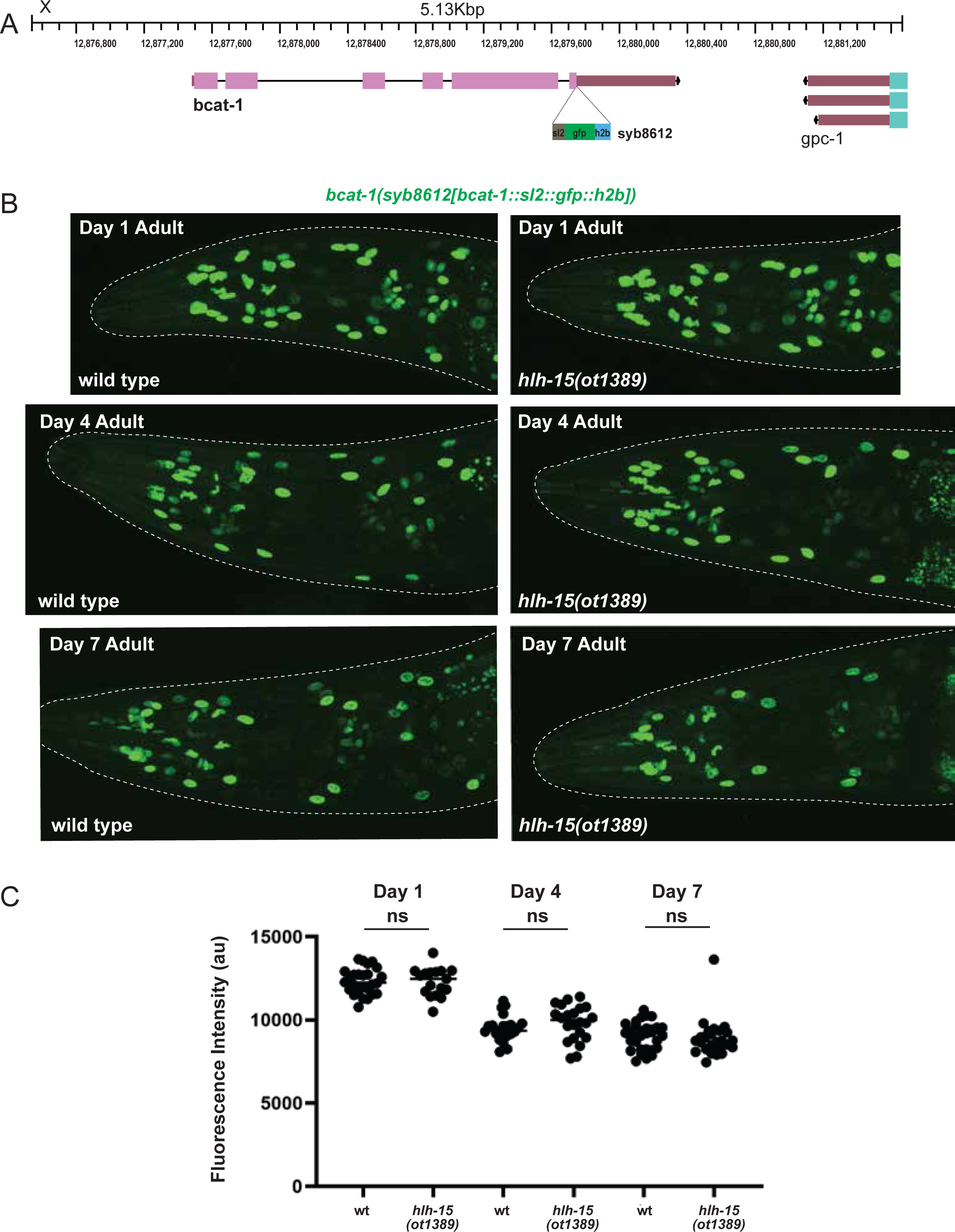
Loss of *hlh-15/NHLH* gene does not affect expression of *bcat-1* reporter allele. **A:** Schematic of *bcat-1* locus showing insertion of *sl2::gfp::h2b* to generate *syb8612* reporter allele. **B:** Representative pictures showing *bcat-1* expression in *hlh-15/NHLH* mutants at day 1, day 4 and day 7 adult worms. Reporter gene used is CRISPR/Cas9-engineered reporter allele *bcat-1*(*syb8612*). **C:** Quantification of *bcat-1(syb8612)* expression in *hlh-15/NHLH* mutants. Animals were scored at day 1, day 4 and day 7 adult stage. Statistical analysis was performed using unpaired T-test.

**Table S1:**
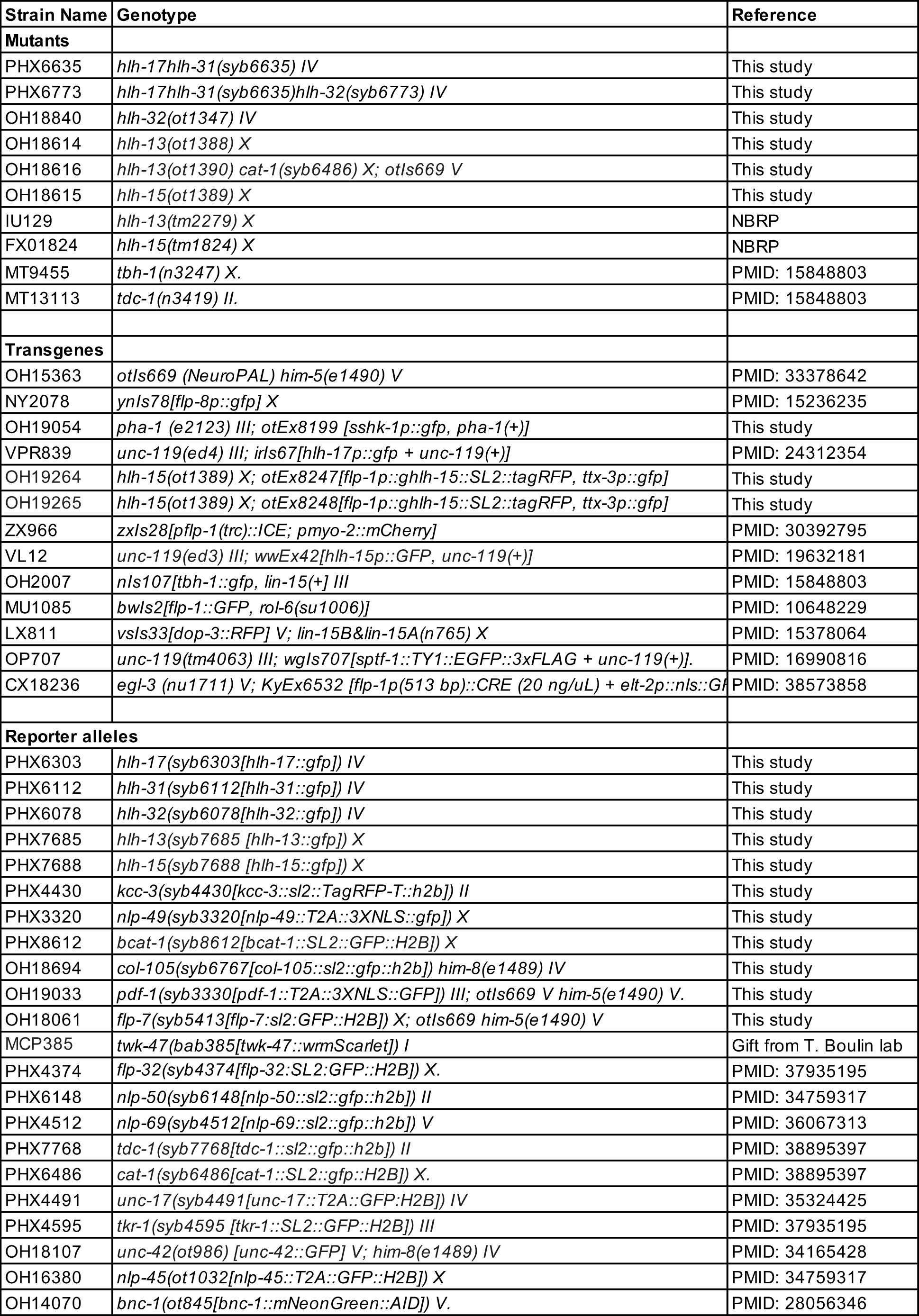
Strain list.

